# Mechanism of small heat shock protein client sequestration and induced polydispersity

**DOI:** 10.1101/2024.12.03.626640

**Authors:** Adam P. Miller, Steve L. Reichow

## Abstract

Small heat shock proteins (sHSPs) act as first responders during cellular stress by recognizing and sequestering destabilized proteins (clients), preventing their aggregation and facilitating downstream refolding or degradation^1–3^. This chaperone function is critically important to proteostasis, conserved across all kingdoms of life, and associated with various protein misfolding diseases in humans^4,5^. Mechanistic insights into how sHSPs sequester destabilized clients have been limited due to the extreme molecular plasticity and client-induced polydispersity of sHSP/client complexes^6–8^. Here, we present high-resolution cryo-EM structures of the sHSP from *Methanocaldococcus jannaschii* (*mj*HSP16.5) in both the apo-state and in an ensemble of client-bound states. The ensemble not only reveals key molecular mechanisms by which sHSPs respond to and sequester client proteins, but also provides insights into the cooperative nature of chaperone-client interactions. Engagement with destabilized client induces a polarization of stability across the *mj*HSP16.5 scaffold, proposed to facilitate higher-order assembly and enhance client sequestration capacity. Some higher-order sHSP oligomers appear to form through simple insertion of dimeric subunits into new geometrical features, while other higher-order states suggest multiple sHSP/client assembly pathways. Together, these results provide long-sought insights into the chaperone function of sHSPs and highlight the relationship between polydispersity and client sequestration under stress conditions.

## INTRODUCTION

Protein aggregation can result from chemical or environmental stressors (*e.g.,* heat or cold, oxidative damage, pH changes) and is a hallmark of many age-related and neurodegenerative diseases such as Alzheimer’s disease (β-amyloid and tau aggregation), Parkinson’s disease (huntingtin aggregation), as well as age-related cataract (lens crystallin aggregation) and cancers^9–16^. Prevention of irreversible protein aggregation in cells is mediated by a proteostasis network comprising chaperones, proteases, and other co-chaperones and regulatory proteins. During cellular stress, small heat shock proteins (sHSPs) act as a front line defense to prevent proteotoxic aggregation by detecting and sequestering partially unfolded, aggregation-prone proteins (aka, clients) through an ATP-independent chaperone ‘holdase’ function^17–19^. During stress recovery, sHSP/client complexes can interact with and deliver clients to ATP-dependent refolding chaperones such as the HSP70 system^3,20,21^. In addition to stress-induced activation or upregulation, mutations in human sHSPs are also associated with multiple protein aggregation diseases, making them intriguing pharmacological targets^9–13^. However, the large and heterogeneous nature of these complexes, along with the promiscuous client interactions typical of sHSPs, has limited mechanistic insights into their ‘holdase’ function.

Many sHSPs possess a high-degree of structural plasticity and undergo mechanisms of client-induced polydispersity characterized by variable sHSP-to-client stoichiometries^7,22^. At the subunit level, sHSPs are relatively small proteins (∼12–43 kDa) with a conserved tripartite domain organization, composed of a central alpha-crystallin domain (ACD) that is well-conserved and flanked by a variable N-terminal domain (NTD) implicated in client-interactions and a flexible C-terminal domain (CTD) involved in sHSP oligomerization^23^. Many sHSPs form dimers that may further assemble into large oligomeric structures (200–800+ kDa) through multivalent intra- and inter-protomer domain interactions. The resulting cage-like sHSP oligomeric assemblies are conserved across all kingdoms of life, along with other morphologies such as disc-like oligomers, fibrillar assemblies, as well as low-order dimer/tetramer forms—showcasing the diversity of the sHSP structural landscape^24–36^.

The first high-resolution structure of an oligomeric sHSP was that of *mj*HSP16.5, from the thermophilic archaeon *Methanocaldococcus jannaschii*, which forms an octahedrally symmetric cage (∼12 nm in diameter) composed of twelve ACD dimers tethered by canonical ACD–CTD interactions^24^. In this structure, the NTD (residues 1–32) implicated in oligomer assembly and client-interaction was completely unresolved, attributed to the flexibility or disorder of this region. However, the structure suggested that the NTD resides inside the caged assembly. Subsequent studies have proposed multiple conformations of the NTD—including α-helical regions—and have resolved a portion of the NTD of *mj*HSP16.5^37,38^. These observations reflect the broader underlying structural plasticity of sHSPs that supports their efficient ‘holdase’ function, that is further mediated by subunit exchange dynamics and oligomeric polydispersity, effectively coupling these structural adaptations to functional demands during stress responses^35,39,40^.

Here, we used single-particle cryo-electron microscopy (cryo-EM) to unveil structural details of both the underlying molecular plasticity of *mj*HSP16.5 in the apo-state, as well as capturing an ensemble of higher-order oligomeric states displaying a variety of configurational features that are induced by the sequestration of heat-destabilized lysozyme, a model client. Structural and mutational analysis, coupled with biophysical and functional characterizations, suggests that multimodal interactions involving the NTD, particularly conserved hydrophobic regions, along with flexible CTD tethering, play critical roles in sHSP assembly, plasticity, and client sequestration. The cryo-EM data suggest a ‘polarized assembly’ mechanism, enabling the sequential recruitment of additional dimer/client units facilitated by a localized destabilization of the sHSP scaffold that is proposed to enable cooperative mechanisms of client sequestration. Together, these findings provide new structural models for sHSP-mediated proteostasis under stress conditions, shedding a critical light on how these complexes could be targeted in diseases associated with protein misfolding.

## RESULTS AND DISCUSSION

### Structural plasticity of the mjHSP16.5 apo-state and the NTD

For structural and functional characterization, *mj*HSP16.5 was recombinantly expressed in bacterial cells and purified to homogeneity without genetic tags or other modifications (Extended Data Fig. 1 and Methods). To assess structural effects of temperature-induced activation of thermophilic *mj*HSP16.5, the purified apo-state oligomers were incubated at either 37° C (apo-37, inactive state) or 75° C (apo-75, activated state) prior to vitrification for cryo-EM studies. Both conditions yielded consensus 3D reconstructions with similar overall features, exhibiting a ∼12 nm cage-like assembly of 24 subunits, consistent with previous studies^24,37,41,42^ (Fig. 1a-e and Extended Data Fig. 2-4).

**Figure 1.**
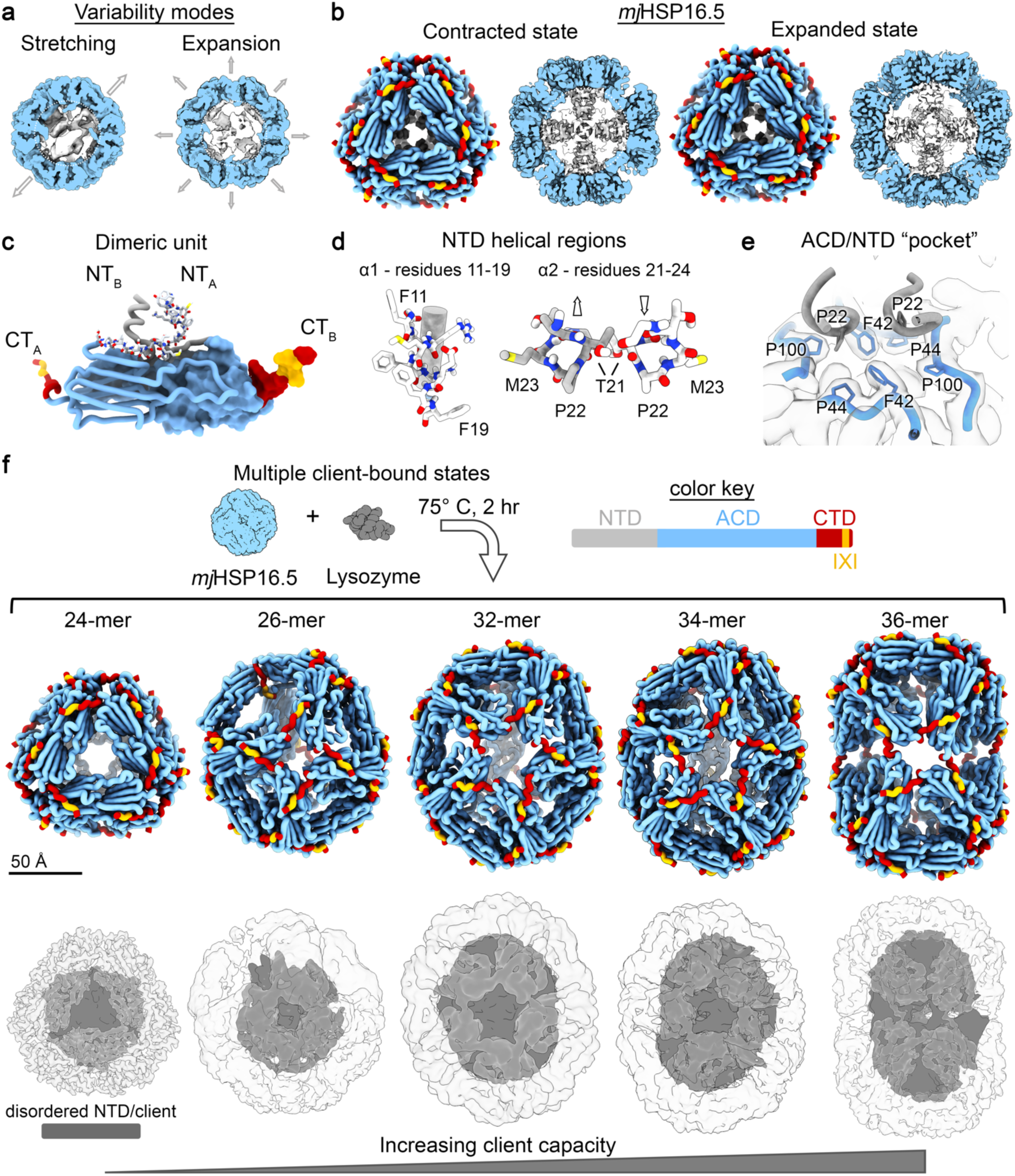
Single-particle cryo-EM analysis of *mj*HSP16.5 in the absence and presence of destabilized client. **a**, *mj*HSP16.5 apo-state displayed conformational dynamics, described by principal component modes of stretching (left) and expansion (right) identified by cryo-EM 3D variability analysis. Asymmetric cryo-EM density maps are displayed in slice view to show the central cavity, with ACDs colored in blue and internal density belonging to the NTD colored in gray. **b**, Resolved contracted (left) and expanded (right) states of *mj*HSP16.5 apo-state (37° C). (left) Atomic models depicted in cartoon representation (ACD:blue, CTD:red, CT-IXI motif:yellow). (right) Central slice of the corresponding cryo-EM density map, colored as in (a). **c**, sHSP dimeric unit of the expanded state with NTD region colored as in (b) and throughout Fig.1 (see color key). **d**, Atomic models of NTD helical regions *α*1 (residues 11-19, left) and *α*2 (residues 21-24, right) with various residues labeled for orientation purposes**. e**, Atomic model and cryo-EM map (semi-transparent) showing the ACD/NTD pocket pertaining to ACD residues (Phe42, Pro44, and Pro100) and Pro22 region of the NTD that fits into the ACD ‘pocket’. **f**, Atomic models (top) and cryo-EM density maps (bottom) of the 24-mer, 26-mer, 32-mer, 34-mer, and 36-mer states of *mj*HSP16.5 obtained in the presence of destabilized lysozyme (75° C incubation for 2 hours). Cryo-EM density maps for the ACD/CTD scaffold shown in transparency, with internal density corresponding to disordered NTD/client shown in gray.

The twelve dimeric building blocks exhibit the canonical β5-β7 loop exchange architecture characteristic of non-metazoan sHSPs, along with ACD-groove/CTD-IXI tethering interactions between neighboring dimers, supporting an overall octahedral symmetry. Asymmetric reconstructions revealed a mix of disordered and helical densities within the cage cavity and lining the inner ACD surface attributed to the NTD. Using multi-class *ab initio* model generation and heterogeneous 3D classification, we identified two similar yet distinct cage morphologies in the apo-37 dataset. Three-dimensional variability analysis (3DVA) revealed principal component modes indicating stretching and expansion of the outer cage-like scaffolding, accompanied by rearrangement of the NTD (Fig. 1a and Extended Data Fig. 3). Further classification and refinement with octahedral symmetry yielded a contracted-state at 2.50 Å resolution and an expanded-state at 2.35 Å resolution (Extended Data Fig. 3; Extended Data Table 1; Supplemental Movies 1 and 2). For both states, strong NTD density was resolved inside the cage and lining the inner ACD surface (Fig. 1b). The apo-75 dataset showed a similar octahedral 24-meric caged assembly resolved at 2.86 Å resolution, with NTD density buried within the cage interior and displayed similar stretching and expansion variability modes (Extended Data Fig. 4).

Atomic models were constructed for each of these states, encompassing the complete ACD, CTD, and most of the NTD (residues 11–32) (Fig. 1b-e; Extended Data Fig. 3-4; Extended Data Table 1; Supplemental Movie 1). Although some differences in the NTDs were observed among the apo-state models (Fig. 2b-d), the overall topology is shared, consisting of a helix-turn-helix-turn-β-sheet structure, denoted as α1 (residues 11–19), α2 (residues 21–24), and β0 (residues 30–32). The NTD forms extensive inter- and intra-domain interactions. Dimeric building blocks exhibit domain-swapped interactions involving α1, with stabilizing salt bridges formed between charged residues Glu12 and Lys16 (Fig. 1c,d). The semi-helical α2 region exhibits intra-dimer hydrogen-bonding potential between Thr21 residues of opposing subunits (Cβ distance 3.1 Å in the apo-37 expanded state) (Fig. 1d). Pro22 is adjacent (< 5 Å) to the inner surface of the ACD, lining an “ACD/NTD pocket” formed by Phe42, Pro44, and Pro100 at the ACD dimer interface (Fig. 1e). Additional stabilizing Van der Waals contacts occur between Met28 (chain A) and Phe19 (chain B) (Cγ distance 4.5 Å in the apo-37 expanded state, Extended Data Fig. 3).

**Figure 2.**
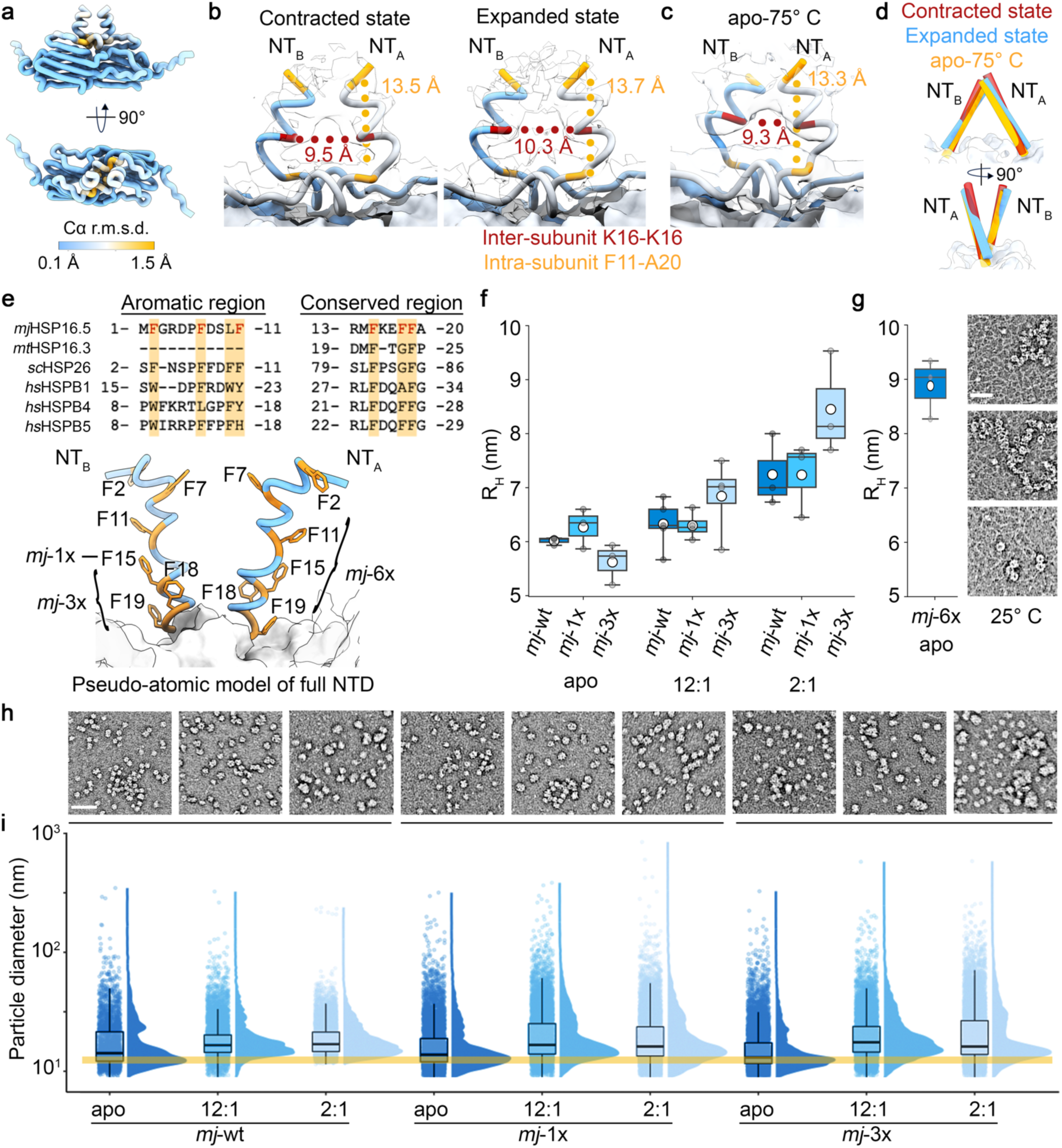
Conserved Phe residues within the NTD are critical to sHSP assembly, stability and chaperone function. **a**, Cα root-mean square standard deviation (r.m.s.d., colored) comparison between the contracted and expanded states of the apo-37 dataset. **b-c**, Distance measurements between NTDs from neighboring chains within a dimer (NT_A_ and NT_B_) for the contracted and expanded states of apo-37 (b) and apo-75 (c). Measurements made between C*α* atoms of Lys16 of each chain, and between Phe11 with Ala20 within a chain. **d**, Views of α1 helical region of the contracted (red) and expanded (blue) states of apo-37 and apo-75 (yellow). **e**, Sequence alignment (top) of ‘aromatic’ and ‘conserved’ regions within the NTD of various small heat shock proteins (*mj*HSP16.5, *M. tuberculosis mt*HSP16.3, *S. cerevisiae sc*HSP26, and human *hs*HSPB1 (HSP27), *hs*HSPB4 (αA-crystallin), and *hs*HSPB5 (αB-crystallin). Phe residues of *mj*HSP16.5 mutated in this study are highlighted. Pseudo-atomic model (bottom) of full *mj*HSP16.5 NTDs within a dimer (NT_A_ and NT_B_) with each Phe residue labeled. **f**, Hydrodynamic radii of major populations detected by DLS for *mj*HSP16.5 wildtype (*mj*-wt) and NTD variants (F15A;*mj*-1x and F15/18/19A;*mj*-3x) in the absence (apo) or presence of lysozyme (12:1 and 2:1, chaperone:client ratios) after incubation at 75° C for 2 hours (n = 3–5 independent experiments). **g**, Hydrodynamic radius (n = 3 independent experiments) and representative NS-EM micrographs of the *mj*-6x variant (at 25° C) in the apo-state (scale bar = 50 nm). **h-i**, NS-EM micrographs (scale bar = 50 nm) and associated Feret diameters obtained from single-particle measurements for *mj*-wt and the *mj*-1x and *mj*-3x variants in the absence (apo) and presence (12:1 and 2:1) of lysozyme after incubation at 75°C for 2 hours displayed as rain cloud plots. Yellow bar indicates the Feret diameter for the major population of particles in the *mj*-wt apo-state dataset. Box plots (panels f,g) show the central 50% of the data, spanning from the first quartile (Q1) to the third quartile (Q3), with a line at the median. Whiskers extend to the minimum and maximum values.

At the subunit level, the expanded and contracted states from the apo-37 dataset are highly similar over the ACD and CTD domains (Cα root-mean-square deviation (r.m.s.d.) = 0.3 Å; Fig. 2a), with the most significant variation localized to the NTD (Cα r.m.s.d. reaching 1.5 Å). The domain-swapped α1 segment maintains regular helical structure in both states, but intra-molecular distances are displaced ∼0.8 Å in the expanded state compared to the contracted state (Fig. 2b). The apo-75 consensus model is also similar to the apo-37 states but exhibits a distinct NTD arrangement (Fig. 2c,d). The α1 separation at Lys16 of apo-75 is ∼9.3 Å versus 9.5–10.3 for the contracted and expanded apo-75 states, respectively (Fig. 2c) The density for the α1 segment, particularly around Phe18 and Phe19, is notably weaker in the apo-75 model, likely reflecting enhanced flexibility at this site (Extended Data Fig. 4).

Assessment of asymmetric 3D reconstructions showed the α1 domain-swap and ‘ACD/NTD pocket’ interactions are uniformly adopted throughout the oligomer. However, 3D reconstructions extracted from 3DVA identified additional asymmetric NTD interactions involving the unmodeled distal region (residues 1– 10). These interactions are dynamically remodeled within the inner cavity during oligomer expansion and contraction, suggesting multiple transient interactions between NTDs (Extended Data Fig. 3). Together, these NTD interactions appear to support both dimer stability and higher-order oligomer assembly, further supporting the notion that sHSP NTDs are highly dynamic and facilitate sHSP oligomer plasticity. This concept is reinforced by the structure of the plant HSP16.9, where 6 of 12 NTDs were resolved, which also implicate this domain in facilitating long-range stabilization across the oligomeric structure^25^.

Interestingly, temperature-dependent activation of *mj*HSP16.5 at 75° C resulted in relatively minor structural rearrangements compared to the apo-37 models, in contrast to other higher-order sHSPs that undergo gross morphological changes in response to temperature^7,40,43^. This suggests *mj*HSP16.5 activation at higher temperatures is primarily facilitated by enhanced subunit exchange dynamics^44^, which were too transient to be captured in our cryo-EM dataset.

### Conserved phenylalanine-rich regions of the NTD mediate oligomeric assembly and stability

The distal NTD region of *mj*HSP16.5 (residues 1-11) forms the so-called ‘aromatic region’, rich in Phe residues. The α1 motif corresponds to the ‘conserved region’, where several Phe residues are also localized and spaced 3-4 residues apart. These Phe residues (F2, F7, F11, and F15, F18, F19) are highly conserved across sHSPs from diverse species, including humans (Fig. 2e). It has been proposed that these Phe-rich regions provide a favorable environment for non-specific transient interactions, enhancing heat stability and supporting the structural plasticity necessary for chaperone function^45,46^.

To interrogate the role of these conserved Phe residues, three variants were generated to replace Phe with Ala: F15A (*mj*-1x), F15/18/19A (*mj*-3x), and F2/7/11/15/18/19A (*mj*-6x). The F15A variant (*mj*-1x) targets the middle of the NTD α1 helical region, positioned between potential π-stacking partners F11, F18, and F19. The *mj*-3x and *mj*-6x variants eliminate Phe residues throughout the ‘conserved’ region and all NTD Phe residues, including both the unresolved ‘aromatic’ region and the α1 ‘conserved’ region, respectively (Fig. 2e). The *mj*-1x and *mj*-3x variants were purified in high-yields; however, the *mj*-6x variant exhibited diminished solubility and stability, resulting in low yields and limiting some analysis. Deletion constructs truncating the NTD at positions 20 and 32 (*mj*-NTDΔ20 and *mj*-NTDΔ32) were also attempted but did not yield soluble protein during expression and were therefore not characterized.

Structurally, *mj*-1x and *mj*-3x variants formed oligomers approximately the same size as wildtype, as determined by dynamic light scattering (DLS) at 75° C. The measured hydrodynamic radii (R_h_, average ± sem with n independent experiments) were 6.3 ± 0.2 nm (n = 3) and 5.6 ± 0.2 nm (n = 3), respectively, compared to 6.0 ± 0.04 nm for the wild type (n = 3) (p = 0.18 and 0.11, respectively) (Fig. 2f). In contrast, *mj*-6x exhibited a significantly larger R_h_ of 8.9 ± 0.6 nm (n = 3; p < 0.01; Fig. 2g) compared to *mj*-wt. Of note, *mj*-6x R_h_ values were measured at 25° C due to the diminished heat stability, showing an aggregation temperature (T_agg_) ∼60° C. In comparison, *mj*-1x and *mj*-3x remained soluble up to 85° C, the highest temperature tested (Extended Data Fig. 1).

To gain more detailed morphological insights, negative stain EM (NS-EM) was performed on each of the variants (Fig. 2g,h and Extended Data Fig. 1). Individual complexes were analyzed directly from raw micrographs by extracting Feret diameters (D_Feret_), describing the largest diameter of a particle^8^ (Fig. 2i). The primary modes in the distribution of D_Feret_ measurements agree with R_h_ values measured by DLS, corresponding to 11.9 nm for wildtype apo-state (wt-apo), 12.2 nm for *mj*-1x, and 10.7 nm for *mj*-3x. Additionally, significantly different degrees of polydispersity related to minor populations of larger complexes or clusters were detected (p < 0.0005, K-S test, Fig. 2i). The *mj*-1x and *mj*-3x particles exhibited the canonical spherical cage-like morphology of wt-apo, appearing as isolated particles and clustered oligomers. Oligomer clusters formed by *mj*-wt, *mj*-1x, and *mj*-3x reached sizes greater than 100 nm, while the constituent oligomers appeared to retain normal morphology, suggesting minimal disruption to the cage-like structure upon clustering (Fig. 2h). Intriguingly, *mj*-6x displayed ring-like structures with diameter of 21.3 ± 2.3 nm (average ± sem), as well as smaller spherical particles and thin filamentous structures approximately 2 nm wide that formed extensive networks (Fig. 2g, *inset* and Extended Data Fig. 1), indicating a substantial impact on the native quaternary structure.

These results suggest the conserved Phe regions within the intrinsically disordered ‘aromatic region’ and ‘conserved region’ are critical for mediating canonical 24-mer assembly and oligomeric stability. The Phe-rich character of the ‘aromatic’ and ‘conserved’ regions in the NTD of sHSPs may support oligomeric condensation through the hydrophobic-effect, while facilitating oligomer plasticity through formation of multiple quasi-equivalent interactions. This is consistent with previous studies that show truncation of the ‘aromatic region’ of *mj*HSP16.5 resulted in smaller oligomers^47^, and mutation of the ‘conserved region’ of human *α*-crystallins (HSPB4 and HSPB5) alters oligomeric size, thermal stability, and chaperone activity^48,49^.

### Client-induced polydispersity of mjHSP16.5

To investigate the chaperone function of *mj*HSP16.5, we established binding conditions that mimic early cellular stress conditions, where the client protein is destabilized (enhanced sampling partially unfolded states) but not yet aggregating. Hen egg lysozyme (14.3 kDa) was chosen as a suitable binding partner because its melting temperature (T_m_) of approximately 75° C is well above the activation temperature of *mj*HSP16.5 (∼60° C), necessary for initiating the subunit exchange dynamics required for chaperone function^44,50^. As controls, *mj*HSP16.5 showed no chaperone protection against reduction-induced aggregation of lysozyme at 37° C, and lysozyme exhibited no observable heat-induced aggregation by DLS up to ∼80° C (Extended Data Fig. 5). Binding assays were performed at chaperone-to-client ratios of 12:1 and 2:1, with *mj*HSP16.5 (wildtype and variants) with the client lysozyme held at 10 µM. Mixed components were incubated at 75° C for 2 hours (*i.e.,* near the T_m_ of lysozyme but below T_agg_), and R_h_ was monitored by DLS.

Wildtype chaperone/client (*mj*-wt/lyso) complexes displayed increased R_H_ values by DLS with exhibited expanded and elongated morphological features visualized by NS-EM, compared to the apo-state, that were more pronounced at the more saturating 2:1 chaperone-to-client ratio (Fig. 2f and Extended Data Fig. 5). The *mj*-1x/lyso and *mj*-3x/lyso 12:1 reactions yielded complexes of similar size to *mj*-wt/lyso, with R_h_ values of 6.2 ± 0.2 nm for *mj*-wt/lyso (n=4), 6.3 ± 0.2 nm for *mj*-1x/lyso (n=5), and 6.8 ± 0.4 for *mj*-3x/lyso (n=4) measured by DLS (Fig. 2f and Extended Data Fig. 5). At the more saturating 2:1 ratio, the formed complexes all displayed larger R_h_ values, including for *mj*-wt/lyso complexes (7.2 ± 0.4, n=3), *mj*-1x/lyso (7.2 ± 0.4 nm, n=3) and *mj*-3x/lyso (8.5 ± 0.6 nm, n=3), although the differences between these variants were not significant (p = 0.49 and 0.07, respectively). Stable complex formation was confirmed for all reactions by size-exclusion chromatography (SEC) and SDS-PAGE, which also demonstrated increased size and polydispersity following complex formation with lysozyme (Extended Data Fig. 5).

Single-particle size analysis by NS-EM allowed for direct visualization and more detailed quantitative analysis of the size and polydispersity of chaperone complexes induced by the sequestered client (Fig. 2h,i and Extended Data Fig. 6). For *mj*-wt the primary D_Feret_ mode increased from 11.9 nm in the apo-state to 13.9 nm and 14.4 nm at 12:1 and 2:1 chaperone:client ratios, respectively, consistent with R_h_ values by DLS. The *mj*-1x and *mj*-3x variants exhibited similar increases in D_Feret_ upon lysozyme binding, reaching 14.5 nm for *mj*-1x at both ratios, and 14.0 nm and 14.5 nm for *mj*-3x at 12:1 and 2:1 ratios, respectively. In all cases, the degree of polydispersity, reflected in the variation of size distributions for the lysozyme-bound complexes, was significantly different from that of the apo-states (p < 0.0005, KS test). Morphologically, client-bound states of *mj*-wt, *mj*-1x, and *mj*-3x exhibited expansion, elongation, and oligomer clustering, where clusters formed by *mj*-3x appeared larger and more amorphous, potentially reflecting the loss of stable NTD interactions.

### Cryo-EM resolved structures of polydispersed chaperone/client complexes

We next sought to characterize the client-bound states for wildtype *mj*HSP16.5 formed with lysozyme at a 12:1 chaperone-to-client ratio using cryo-EM. Under these conditions, the formed complexes were closer in size to the apo-state, suggesting a more tractable target for high-resolution analysis. After incubating the mixture at 75° C for 2 hours, the specimen was vitrified and subjected to cryo-EM data collection and single-particle analysis (Extended Data Fig. 7). Two-dimensional classification and multi-class *ab initio* model generation revealed the presence of a range of client-induced *mj*HSP16.5 oligomeric states (Extended Data Fig. 7). Ultimately, five different structures were resolved at sufficient resolution to model the ACD and CTD regions, corresponding to the octahedral 24-mer (2.6 Å), asymmetric assemblies of a 26-mer (3.7 Å), 32-mer (4.4 Å) and 34-mer (4.7 Å), as well as a symmetric 36-mer (4.0 Å, D3-symmetry) (Fig. 1f; Extended Data Fig. 7; Supplemental Movie 3). Notably, there was no evidence of monomer insertion/removal between the resolved oligomers, consistent with the dimer functioning as the primary unit of the *mj*HSP16.5 chaperone.

These client-bound states showcase the intrinsic ability of sHSPs to form diverse oligomeric assemblies using fundamental building blocks, where coupling through the flexible CTD facilitates formation of structurally distinct features (windows and axes) compared to the canonical 24-mer. The internal volume of the cage assemblies increases from the 24-mer to the 36-mer, presumably reflecting enhanced client sequestration capacity in higher-order oligomeric states (Fig. 1f). In all client-bound states, the internal density exhibited lower local resolution with unresolvable features, consistent with either extensive client unfolding in the sHSP-bound state or multiple transient binding interactions that prevented alignment for 3D reconstruction.

The resolved 24-mer under these conditions closely resembled the apo-states, with comparable NTD densities and stretching/expansion modes of variability (Extended Data Fig. 7). Notably, the internal cavity of the 24-mer is nearly filled with density belonging to the NTD (∼19 nm^3^ of non-NTD density), and thus appears incapable of sequestering clients as large as lysozyme (∼14 kDa, ∼22 nm^3^). It is therefore likely that this state represents an unbound population of *mj*HSP16.5 particles. This notion is further supported by the observation that the NTD in the higher-order oligomers becomes unresolved (Fig. 1f), consistent with an order to disorder transition that accompanies client sequestration, likely due to complex forms of client-interactions. However, even a modest transition to a 26-mer induces a dramatic increase of the internal non-NTD volume to ∼100 nm^3^, sufficient to sequester this client. The concomitant increase in the cavity volume established by the formation of higher-order assemblies (34-mer and 36-mer) would be expected to facilitate higher capacity client binding, enabling a single sHSP complex to sequester multiple clients or clients of variable size. Indeed, the non-NTD volume inside the 36-mer (∼140 nm^3^) would be sufficient for sequestering up to six lysozyme molecules (Fig. 1f).

### CTD flexibility enables client-induced sHSP polydispersity

Superimposing the component subunits from each of the assemblies obtained in the presence of client reveals the flexibility of the CTD, which facilitates the tethering of ACD dimers within the various oligomeric states (Fig. 3a). Cα r.m.s.d. analysis showed minimal variability within the core ACD region across all states, appearing similar to the apo-state. In contrast, the CTD adopts an apparent continuum of states within the various assemblies, with two primary modes of CTD extension (up and down) that are most distinctly separated in the symmetric 36-mer, resulting in ∼10 Å Cα deviation of the CTD IXI motif in these two modes (Fig. 3a, *far right denoted by * and †*).

**Figure 3.**
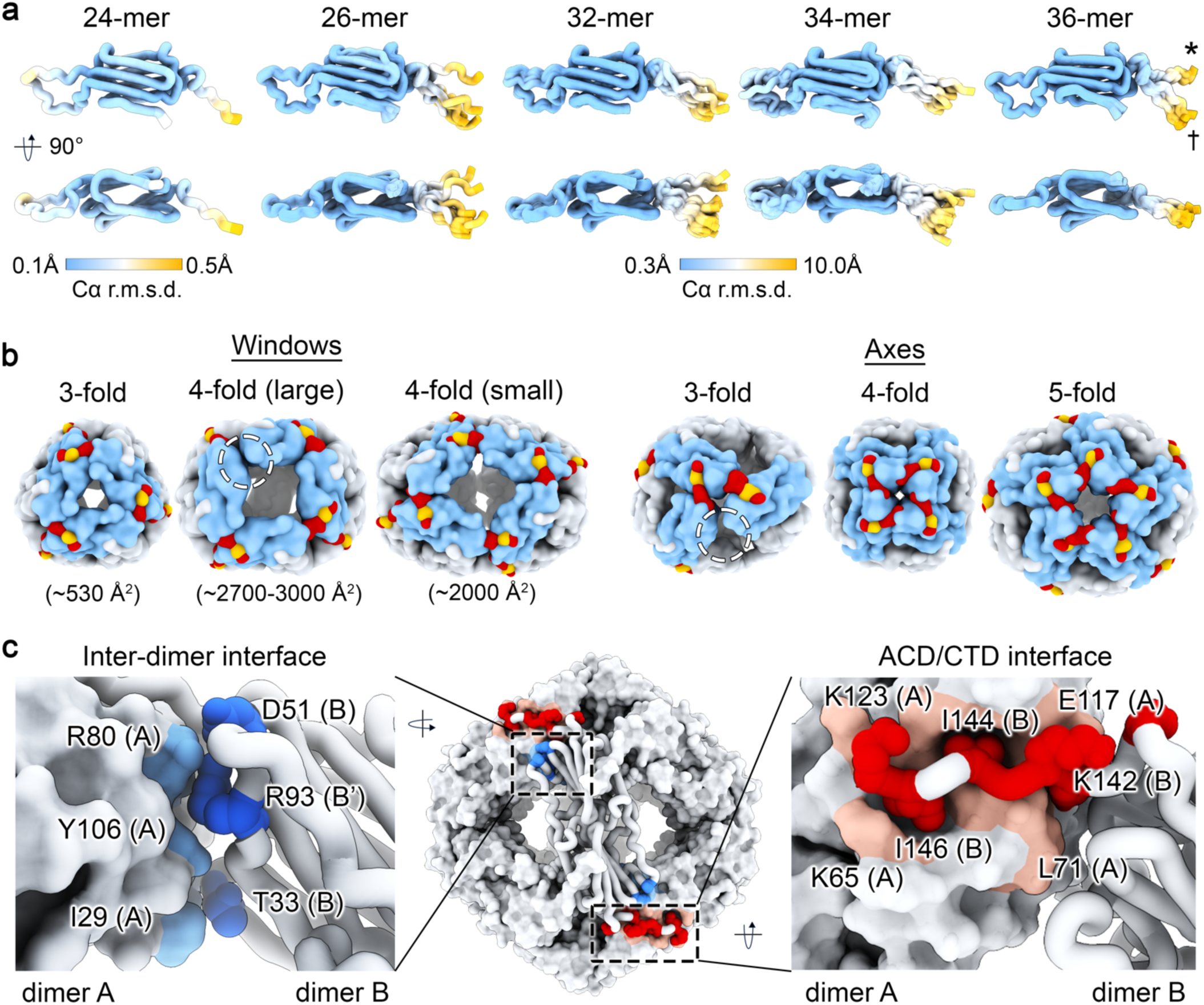
Flexible CTD interactions facilitate multiple oligomeric states that correlate with adopted non-canonical geometrical features. **a**, Comparison of Cα r.m.s.d. values for protomers within each oligomer state obtained in the presence of lysozyme (24-mer, 26-mer, 32-mer, 34-mer, and 36-mer shown left to right). Color keys show range of r.m.s.d. values for the 24-mer (left) and other higher-order oligomers (center). **b**, Windows and axes adopted by the 24-mer (canonical 3-fold window/4-fold axis), and non-canonical states formed by the 26-mer (4-fold window/3-fold axis), 32- and 34-mer (4-fold window/5-fold axis), and 36-mer (4-fold window/4-fold axis). sHSP models displayed as low-pass filtered surfaces with subunits colored to highlight the specified features. Circled areas highlight unresolved/unmodeled CTD regions, presumably reflecting intrinsic flexibility. **c**, Interfaces corresponding to the inter-dimer interfaces formed between ACDs (blue/left) and ACD/CTD interaction (red, right). Zoom views, depict residues involved in these interfaces.

The flexible CTD interactions facilitate the formation of novel structural features in higher-order oligomeric assemblies of *mj*HSP16.5. The apo-state 24-mer forms eight canonical 3-fold windows (∼530 Å^2^) and six 4-fold axes (Fig. 3b). In comparison, the 26-mer establishes two large 4-fold windows (∼3000 Å²) and two 3-fold axes; the 32-mer and 34-mer also display large 4-fold windows (∼2700 Å², two and one each, respectively) and 5-fold axes (one and two each, respectively); and the 36-mer exhibits three smaller 4-fold windows (∼2000 Å²) and six 4-fold axes (Fig. 3b). The increased exposure of the internal cavity along the expansive 4-fold windows could provide fenestrations that enable NTDs from within the cage to form hydrophobic interactions with exchanging or unbound NTDs/clients. Indeed, previous studies have shown that larger complexes formed by engineered variants of *mj*HSP16.5 exhibit increased affinity toward clientele, proposed to be attributed to increased hydrophobic surface area^51,52^.

The conformations of the CTD correlate with these new geometric features of the caged assemblies. Specifically, the canonical 3-fold windows and newly identified 5-fold axes are facilitated by the CTD in the downward conformation (Fig. 3a, *†)*, whereas the 4-fold windows in the higher-order states require an upward CTD conformation (Fig. 3a, *\**). The 26-, 32-, and 34-mer structures all exhibit weak or absent density corresponding to the CTD region of some monomers and were not modeled, presumably reflecting intrinsic dynamics at these sites (Fig. 3b, *circled*). The dimer connecting two 3-fold axes of the 26-mer was of considerably low local resolution (∼10 Å) and exhibited weak density throughout the ACD region, specifically for the *β*5-*β*7 loop. Similarly, *β*5-*β*7 loop regions in the 32-mer and 34-mer show weak density in the cryo-EM maps and display higher Cα r.m.s.d.’s in the corresponding models (Fig. 3a).

The expanded and contracted models of apo-state 24-mer exhibit inter-dimer interfaces involving the NTDs with surface areas of 133 ± 0.2 Å^2^ and 112 ± 0.3 Å^2^, respectively. The inter-dimer interface is comprised of contacts between the proximal NTDs (residues 29, 30, 33, and 36), *β*5 and *β*7 (residues 78, 80, and 106) with the *β*5-*β*7 loop (residue 93) and establishes the corners of 3-fold and 4-fold cage windows (Fig. 3c, *left*). The canonical ACD/CTD tethering interface of the expanded and contracted 24-mer models establish a contact surface area of 644 ± 0.2 Å^2^ and 669 ± 0.1 Å^2^, respectively. This interface is characterized by tongue-and-groove interactions formed between the CTD-IXI motif of one protomer (residues 138, 141-144, 146, and 147) and the *β*4/8-groove on the ACD of a neighboring protomer (residues 65, 70-73, 76, 78, 80, 117, and 120-123) (Fig. 3c, *right*). Surprisingly, despite the profound display of geometric reconfigurations in the higher-order client-bound oligomers, there was minimal disruption to either the inter-dimer or the ACD/CTD tethering interfaces, with standard deviations of 12 Å^2^ (9.5 %) and 78 Å^2^ (13 %), respectively, across all oligomeric states. Together, these observations mechanistically rationalize how sHSPs can achieve such high-degree of polydispersity from the modular ACD dimer building blocks.

### Client-induced sHSP polarized destabilization directs chaperone elongation

To gain further insights into the regional stability of the various assembly states, local resolution analysis was performed on cryo-EM maps generated from asymmetric 3D reconstructions (Fig. 4a). The 24-mer reconstruction obtained in the presence of lysozyme showed minimal variation in resolution over the exterior scaffold, composed by the ACD and CTD, with local resolution of ∼2.5–3.0 Å. The internal NTD (or potentially NTD–client) density exhibited lower resolution (∼4.5 Å) with similar features to that of the apo-state, including symmetric NTD helices and unassigned NTD elements. In contrast, the intrinsically asymmetric 26-mer, 32-mer, and 34-mer assemblies showed extensive resolution variation across the caged scaffold, and a loss of defined NTD features. Their most well-defined regions exhibited relatively high local resolutions (∼3–4 Å), centered around a single canonical 4-fold axis. Moving away from this vertex along the axis of the elongated cage morphologies, the local resolution continuously decreased, reaching ∼9–10 Å. The 36-mer reconstruction had an overall lower resolution, likely due in part to the limited number of particles in this class, yet this map displayed a more uniform resolution distribution across the elongated scaffold (∼7–8 Å) reflecting the internal symmetry of this assembly state.

**Figure 4.**
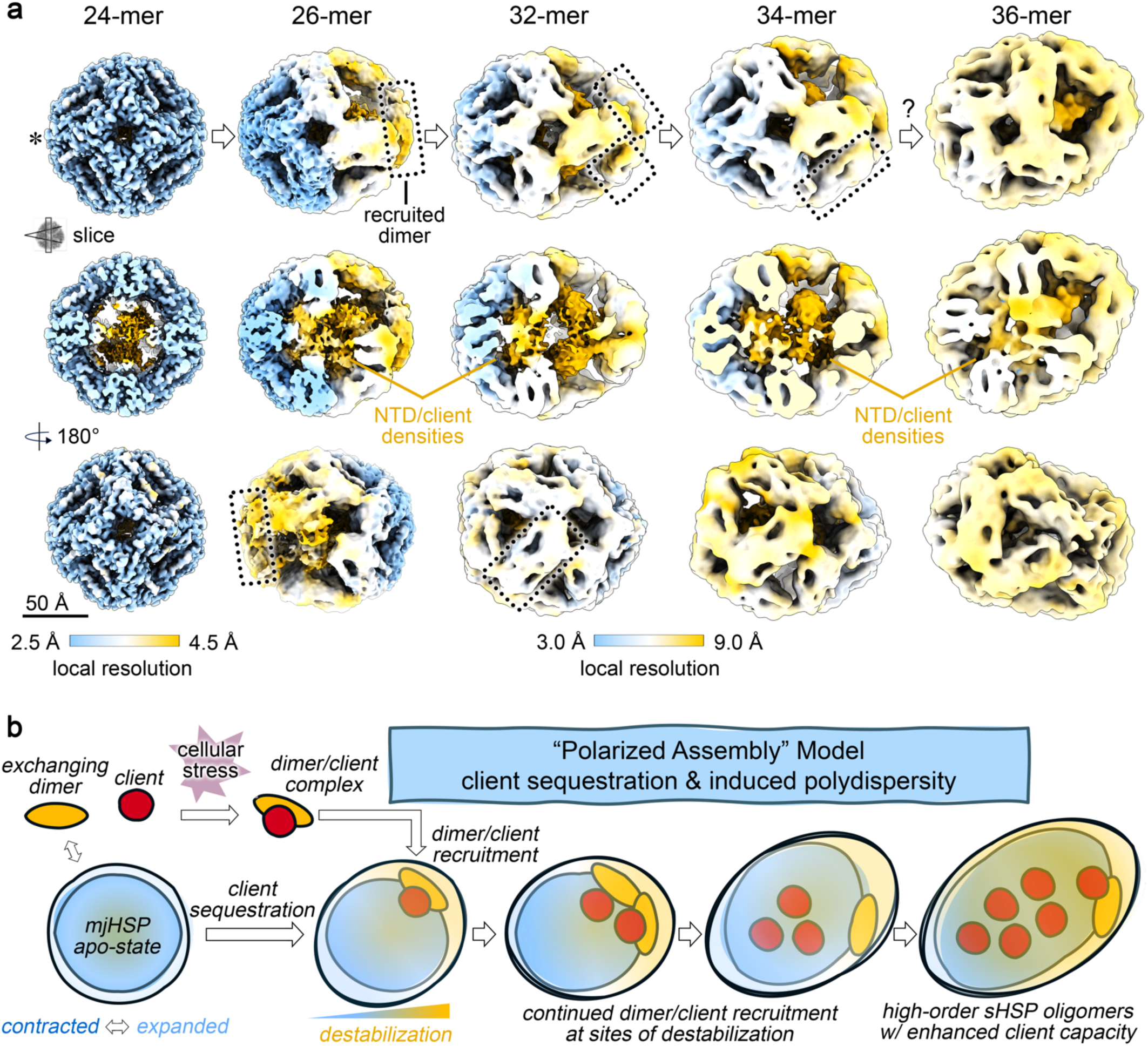
Polarized destabilization induced by client sequestration correlates with sites of recruited dimers to form higher-order sHSP oligomers. **a,** Asymmetric cryo-EM density maps of *mj*HSP16.5/lysozyme oligomeric state depicted along a 4-fold axis (top), an internal slice view (middle) showing NTD/client densities, and a 180° rotation (bottom). Map densities colored according to local resolution. Color keys show range of resolution values for the 24-mer (left) and the other higher-order oligomers (center). Asterisk indicates site of common axis for alignment. Dotted rectangles indicate sites of dimer recruitment to form higher-order oligomers. Scale bar = 50 Å. **b,** Illustration summarizing the proposed “polarized assembly” model of sHSP client sequestration and induced polydispersity, as described in the main text.

By aligning the most ordered regions of the caged assemblies, we can attempt to infer aspects of the chaperone/client assembly pathway based on shared geometrical features between complexes, the polarized nature of local resolution, and variability analysis at the subunit interfaces (Fig. 4a, *outlined dimers*). Notably, the initial step of higher-order oligomeric assembly remains elusive because the apo-state 24-mer would need to undergo significant structural deformation of a canonical 4-fold axis to accommodate the insertion of an additional dimer (or dimer/client complex). Our cryo-EM variability analyses of the 24-mer and 26-mer did not detect any conformational states showing such dramatic rearrangements, possibly due to the transient nature of this proposed transition state. Once initial client sequestration occurs, inducing formation of the 26-mer, it creates two larger 4-fold windows and two destabilized 3-fold axes into which new dimers (or dimer/client complexes) can be inserted or “recruited” onto the complex. Variability analysis of the 26-mer revealed extensive dynamics at the site of the recruited dimer, with a high degree of flexibility between the two new 3-fold axes (Supplemental Movie 4).

Following formation of the 26-mer, further subunit and client recruitment may be facilitated by the polarized destabilization of the sHSP scaffold. This destabilized region would likely promote subunit exchange and direct the formation of higher-order states that can sequester more client proteins (Fig. 4b). This proposed ‘polarized assembly’ model would lead to an effective elongation of the sHSP scaffold, similar to observations in human α-crystallins, as well as murine HSP25 and yeast HSP26^8,53^. The concept of polarized sHSP stability contributing to chaperone activation has also been suggested in the context of HSP26 phosphorylation^54^. Such localized destabilization may be critical for the chaperone mechanism by providing “hot spots” for cooperative subunit recruitment and client insertion enhancing client capacity, or by enabling access and/or reversible release of clients to downstream refolding machinery^55^.

Notably, we did not identify any *bona fide* 28-mer or 30-mer populations in the cryo-EM dataset, suggesting these states either do not form or are intrinsically unstable. A plausible transition from a 32-mer to a 34-mer was uncovered by 3DVA, suggesting that a simple insertion, or recruitment, of a dimer into a large 4-fold window of the 32-mer could readily occur, resulting in the formation of two 3-fold windows and an additional 5-fold axis (Supplemental Movie 4).

While this ‘polarized assembly’ model may account for many of these observations, it also raises several questions. Notably, there appears to be no clear pathway to transition directly from the 34-mer to the 36-mer through simple dimer insertion. Such a transition would apparently require disassembling two 5-fold axes and rearranging multiple dimers to accommodate an additional dimer or dimer/client unit. Therefore, it is possible that the 36-mer, and other sHSP/client states, assemble via an alternative pathway(s). For example, initial complexes formed between a destabilized client and exchanging sHSP dimers could seed *de novo* oligomeric assembly by recruiting other exchanging dimers or dimer/client complexes to form higher-order assemblies. In this scenario, the most energetically favorable sHSP scaffold architecture may be determined by client characteristics, such as aggregation state/size and degree of unfolding, that seeded the interaction.

This alternative “seeded assembly” model is supported by recent single-molecule fluorescence studies on human αB-crystallin and HSP27, which showed that initial binding is followed by additional subunit recruitment, with variations between the two sHSPs indicating chaperone-specific mechanisms^22,56^. Notably, the “polarized assembly” and “seeded assembly” models are not mutually exclusive and may both contribute to the complexity of sHSP sequestration mechanisms. Future studies aimed at capturing the transition states involved in client sequestration are needed to fully elucidate these mechanistic details. Interestingly, despite the potential for multiple higher-order assembly pathways, the formation of directionally elongated cage-like assemblies appears to be an energetically favored aspect of the chaperone sequestration mechanism.

### Concluding remarks

A major hurdle in understanding sHSP chaperone mechanisms has been the polydispersed nature of sHSP/client complexes. This study demonstrates that *mj*HSP16.5 can be harnessed as a model system to explore sHSP polydispersity in relation to chaperone function, guiding future research on the structure and function of other enigmatic sHSP systems. Future efforts to investigate sHSP/client complexation in the cellular context may provide additional insights for targeting co-aggregation pathways in sHSP-related diseases such as cataract, neurodegenerative disease, cardiomyopathies, and cancer treatment resistance^13,14,57^.

## ACKNOWLEDGEMENTS

We thank Dr. Kirsten Lampi for helpful discussions. We are grateful for instrumentation access and training provided by the staff at the OHSU Multiscale Microscopy Core and Advanced Computing Center, and the Pacific Northwest Cryo-EM Center (supported by NIH Grant R24GM154185) with the assistance of Dr. Janette Myers. The research was funded by NIH grants R01EY030987 and R35GM124779 (to S.L.R.) and fellowship F31EY033226 (to A.P.M.).

## AUTHOR CONTRIBUTIONS

A.P.M. performed the cryo-EM studies, biophysical characterizations, functional experiments, and analyzed these data; S.L.R. and A.P.M. conceived of the study and prepared the manuscript.

## METHODS

### Construction of expression plasmids

The gene sequence of wildtype *mj*HSP16.5 (GENID: 1451140) was codon-optimized for bacterial expression and encoded into a pET23a(+) expression vector and sequences encoding variants within the N-terminal domain of *mj*HSP16.5: F15A (*mj*-1x), F15/18/19A (*mj*-3x), F2/5/11/15/18/19 (*mj*-6x), deletion of residues 1-32 (*mj*-Δ32, with Met1 at position 32), and deletion of residues 1-20 (*mj*-Δ20, with Met1 at position 20) were encoded into the pET21a(+) vector (Genscript). Protein expression constructs used in this study did not include solubility or purification tags. Full plasmid sequencing (Plasmidsaurus, Eugene, OR) confirmed the correct gene sequence, insertion site, and placement of mutations/truncations.

### Expression and purification of mjHSP16.5 wildtype and variants

Wildtype (*mj*-wt) and variant constructs (*mj*-1x, *mj*-3x, *mj*-6x, *mj*-Δ32, and *mj*-Δ20) of *mj*HSP16.5 were expressed in bacteria and purified using the same protocols (modified from Quinlan, *at al.*^41^). Briefly, *E. coli* BL21(DE3) was used as an expression host for all constructs and the growth media (LB) was supplemented with ampicillin (0.5 mM). Cells were grown at 37° C to an optical density (A.U. at 600 nm) of 0.6 – 1.0 and expression was induced with 1 mM isopropyl *β*-D-1-thiogalactosidase (IPTG). Cells were harvested by centrifugation (4,000 r.c.f for 15 min at 4° C) after 3-4 hours post-induction at 37° C. Pelleted cells were resuspended in 20 mM Tris-HCl (pH 8.0), aliquoted, and frozen at -80° C for further use.

For purification of each protein construct, cell suspensions were thawed and supplemented with 1,4-dithioreitol (DTT, 0.5 mM final concentration) and phenylmethylsulfonyl fluoride (PMSF, 0.1 mM final concentration), lysed by sonication on ice (70% amplitude, 6 rounds of 30s on/off), supplemented with additional PMSF (0.2 mM final concentration), and cellular debris cleared by ultracentrifugation at 165,000 r.c.f for 30 minutes at 4° C. The supernatant was supplemented with NaCl (1M final concentration) and 20 mM Tris-HCl (pH 8.0) and incubated in an 80° C water bath for 30 minutes (V = 20 mL). The heated lysate was recovered on ice for 5 minutes and denatured protein was pelleted by ultracentrifugation at 165,000 r.c.f for 30 minutes at 4° C. The supernatant containing the thermo-stable *mj*HSP16.5 was collected, DNase I (∼400 units, Thermo Scientific) was added and incubated for 30 minutes on ice before being filter at 0.22 µm prior to chromatography.

The clarified lysate was applied to a gel filtration chromatography column (S300 resin; Pharmacia) equilibrated with 20 mM Tris-HCl (pH 8.0), 1 mM EDTA and 0.5 mM DTT. Fractions from gel filtration were assessed via SDS-PAGE and fractions containing *mj*HSP16.5 (wildtype or mutants) were pooled and supplemented with DTT (0.5 mM final concentration). The pooled fractions were loaded onto a MonoQ anion exchange column (GE Healthcare) equilibrated with buffer A (20 mM Tris-HCl (pH 8.0), 0.16 mM EDTA, and 1 mM EGTA) and eluted with a NaCl gradient (buffer A with 1 M NaCl). Eluted fractions containing the target *mj*HSP16.5 construct were pooled, concentrated to a volume of ∼2 mL with a 100,000 kDa cutoff spin concentrator (Vivaspin), and loaded onto a Superose 6 size-exclusion chromatography (SEC) column equilibrated with 20 mM HEPES (pH 7.4), 2 mM EDTA, 2 mM EGTA, and 100 mM NaCl. Fractions containing purified *mj*HSP16.5 constructs were pooled, aliquoted and either flash frozen in liquid nitrogen and stored at -80° C for later use or incubated at 37° C (wildtype) or 4° C (mutants) for immediate use. Nucleic acid contamination was assessed by monitoring UV absorbance ratio of 280/260 nm with all purified protein having ratios >1.5, indicating minimal co-purification of nucleic acids. The concentration of purified protein was determined by UV absorbance at 280 nm using the extinction coefficient 8,257 M^-1^cm^-1^ ^58^. The variants *mj*-Δ20 and *mj*-Δ32 resulted in insoluble protein and was not purified for downstream analyses.

### Aggregation assays of reduced lysozyme at 37° C

All aggregation assays were performed in reaction buffer containing 20 mM HEPES (pH 7.4), 2 mM EDTA, 2 mM EGTA, and 100 mM NaCl. Aggregation of hen egg-white lysozyme (Fisher, MS grade) at 37° C was induced with the addition of 2 mM tris(2-carboxyethyl)phosphine (TCEP). Lysozyme (10 µM) aggregation by TCEP was monitored in the presence of 120 µM (12:1 ratio) and 20 µM (2:1 ratio)) or absence of *mj*-wt. Chaperone/client mixtures were allowed to equilibrate at 37° C for 15 minutes prior to the addition of TCEP (2 mM final concentration). Turbidity measurements were monitored by absorption at 360 nm, collected in 384-well plates (Nucleon, flat black) on a Tecan Infinite M NANO+ for 2 hours at 37° C.

### Heat induced aggregation and binding assays with lysozyme at 75° C

Dynamic light scattering (DLS) measurements were performed in an Aurora 384 well plate on a Wyatt DynaPro plate reader III (Wyatt Technology, Santa Barabara, USA) operating with an 830 nm laser and 150° DLS detector angle. All measurements were acquired with five reads and 10s acquisition time in the Dynamics software v7.10.1 (Wyatt). To determine the aggregation temperature of lysozyme, the hydrodynamic radius in solution was monitored using DLS while ramping temperature from 25° to 85° C at 0.91° C/min (n = 3). Due to the small size of lysozyme (∼2 nm radius) the working concentration of 10 µM was not detectable and 100 µM was used to determine the aggregation temperature. Likewise, lysozyme at 50 and 100 µM was monitored by DLS at a constant 75° C for 2 hours to show no immediate (< 1 hour) aggregation at these temperatures.

For binding assays, lysozyme (10 µM) was incubated in the presence and absence of *mj*HSP16.5 wildtype, *mj*-1x, and *mj*-3x at 120 µM (12:1 ratio) and 20 µM (2:1 ratio) at 75° C for 2 hours in 20 mM HEPES (pH 7.4), 2 mM EDTA, 2 mM EGTA, and 100 mM NaCl. Prior to incubation at 75° C the mixed samples were incubated at 25° C for 30 minutes. Additionally, samples of *mj*HSP16.5 were measured by DLS in the absence of lysozyme at 25° C (*mj*-6x), or 37° C, 75° C, or through a temperature ramp from 25–85° C at a rate of 0.49° C min^-1^ (*mj*-wt, *mj*-1x, *mj*-3x: 120 µM). The aggregation temperature of *mj*-6x was determined using a heat ramp of 0.3°C min^-1^ from 25–85° C. Replicates of DLS readings were pooled for downstream analyses (SEC/SDS-PAGE, NS-EM, cryo-EM). The average ± sem was calculated for the hydrodynamic radius of (n = 2–3) technical replicates across (n = 3–5) independent experiments for each sample. Statistical significance was assessed by completing a F-test for variability followed by a Student’s two-sided T-test (equal/unequal variance depending on F-test results).

### Size-exclusion chromatography

Following binding assays performed at 75° C, pooled DLS replicates obtained from binding reactions were loaded (125 µL injection) onto a Superose 6 SEC column equilibrated with 20 mM HEPES (pH 7.4), 2 mM EDTA, 2 mM EGTA, and 100 mM NaCl. Elution peaks were monitored by SDS-PAGE (17.5% acrylamide) and protein bands were visualized by silver staining.

### Negative stain EM and single-particle morphology analysis

Negative stain EM was performed on purified apo-state *mj*HSP16.5 that was incubated at 37° C for approximately 16 hr and then diluted to ∼3 µM in dilution buffer containing 20 mM HEPES (pH 7.4), 100 mM NaCl, 2 mM EDTA, and 2 mM EGTA. Chaperone assay reaction products of *mj*-wt, *mj*-1x, and *mj*-3x in the absence (apo) and presence of lysozyme (12:1 and 2:1 chaperone:client ratios) prepared at 75° C were recovered on ice and diluted to ∼ 3 µM (*mj*HSP16.5 concentration) with dilution buffer. Carbon coated 400 mesh copper EM grid (Ted Pella) were glow discharged at 15 mA for 1 min prior to sample application. For each condition, 3 µL of sample was applied to the grid and excess protein/buffer was blotted with filter paper, washed twice with ultrapure water, stained with freshly prepared (0.75% wt vol^-1^) uranyl formate (SPI-Chem), blotted with filter paper, and dried with laminar air flow. Grids of chaperone reactions were set within 30 minutes following ice recovery to quench subunit exchange dynamics following complex formation at 75° C. Negatively stained specimens were imaged on a 120 kV TEM (Tecnai T12, FEI) equipped with either a 2K x 2K CCD camera (Eagle, FEI) at a nominal magnification of 49,000 and a calibrated pixel size of 4.4 Å pixel^-1^ (*mj*-wt apo, 12:1, 2:1) or an AMT camera (model XR16) using the AMT Image Capture Engine (v602.591j) at a nominal magnification of 30,000 with a calibrated pixel size of 4.0 Å pixel^-1^. Micrographs were collected with a defocus range from 1.5–2.2 µm.

Single-particle morphology analysis was performed as previously described^8^. Briefly, unprocessed micrographs were imported into FIJI^59^ and the scale was set based on the calibrated pixel size of the micrograph. Micrographs were processed using the fast-Fourier transform based bandpass filter with default settings (filter large structures at 40 pixels, filter small structures at 3 pixels, 5% tolerance, auto scale after filtering, saturate image when autoscaling) followed by a maximum filter (radius of 2 pixels) and background subtraction (rolling ball radius of 25-50 pixels). The filtered and background subtracted micrographs were binarized (dark background) and segmentation was optimized using the Remove Outliers tool and erosion/dilation of binary segments tools. Processed micrographs were compared to the raw micrograph during optimization of binary segments. The Analyze Particles tool was used for automated determination of Feret diameter of each segment within a minimum particle area of 50 nm^2^. Feret diameters are presented as raincloud plots generated in R Studio (v4.0.5). Statistical analysis was done in Excel (average ± sem) and Scipy^60^ (Kolmogorov-Smirnov test).

### Cryo-electron microscopy data collection

Prior to vitrification, samples were incubated for >16 hr at 37° C (apo-37C) or for 2 hours at 75° C (reaction products from DLS experiments) in the absence and presence of lysozyme (apo-75C and *mj*:lyso-75C at a 12:1 ratio). 3 µL of each sample (∼1 mg/mL) was applied to a freshly glow discharged (15mA, 1 min) holey carbon copper grid (apo-37C: Cflat (EMS) R1.2/1.3, apo-75C and *mj*:lyso-75C samples: Quantifoil R2/1, 400 mesh). Grids were blotted (1.0–1.5 s) at room temperature and 90% humidity and plunge froze into liquid ethane on a Vitrobot Mark IV (FEI, Thermo Fisher Scientific). Image datasets were collected at the Pacific Northwest Cryo-EM Center (OHSU, Portland, OR) on a 300 kV Titan Krios equipped with a K3 detector (Gatan) using SerialEM^61^. Movies were collected in super-resolution mode at a calibrated physical/super-resolution pixel size of 0.788/0.394 Å pixel^-1^ (apo-37C sample), 1.013/0.506 Å pixel^-1^ (apo-75C sample), and 1.066/0.533 Å pixel^-1^ (*mj*:lyso-75C) with a total dose rate of ∼40 e^-^ per Å^2^ over 70 frames for the apo-37C sample and ∼50 e^-^ per Å^2^ over 50 frames for apo-75C and *mj*:lyso-75C. Movies were collected over a defocus range of 1.0–2.5 µm. The apo-75C and *mj*:lyso-75C samples were collected using a GIF energy filter (Gatan) with a 10 eV slit width.

### Cryo-EM image processing of apo-state mjHSP16.5 (37° C)

All steps of cryo-EM image processing were performed in CryoSPARC v3.3.1^62^. A dataset of 16,214 micrographs for *mj*HSP16.5 (apo-37C) was preprocessed with Patch Motion Correction (micrographs binned 2x, 0.788 Å pixel^-1^) and Patch CTF estimation. Low quality micrographs were removed based on CTF resolution fit. A subset of 100 micrographs was subjected to blob picking (120-160 Å diameter) to yield a particle set of ∼3.4 million particles extracted with binning (2.46 Å pixel^-1^). Noisy particles and low occupancy classes were removed by 2D classification to give a set of 1,170,772 particles used for multi-class *ab initio* model generation with 4 classes and maximum resolution of 6 Å. Multi-class *ab initio* generation yielded 2 good classes (1,060,133 total particles) corresponding to 24-mer caged assemblies with slightly different diameters. Further rounds of 2D classification yielded 968,458 particles that were again subjected multi-class *ab initio* reconstruction (3 classes), yielding two distinct classes of 24-meric cages with 441,253 particles in class 1 (expanded state) and 421,283 particles in class 2 (contracted state). A consensus refinement of re-extracted particles (1.05 Å pixel^-1^) of the combined classes (862,436 particles) without symmetry (C1) yielded a consensus reconstruction at 2.99 Å resolution. 2D classification and heterogeneous refinement (C1, 6 classes) and removal of low occupancy classes yielded 630,757 particles which refined with octahedral (O) symmetry to 2.44 Å.

The C1 consensus refinement of the 630,757 particle stack was used as input for 3D Variability analysis with 3 orthogonal principal modes and a filter resolution of 5 Å^63^. Additionally, this particle set was expanded with octahedral (O) symmetry (15,136,968 expanded particles) and used for 3D Variability analysis with 3 orthogonal principal modes and a filter resolution of 5 Å. These particles were subjected to heterogeneous refinement with four classes which gave two high occupancy classes at ∼2.7 Å resolution (expanded state: 257,168 particles; contracted state: 205,181 particles). Separate cleanup of the two particle sets was done by 2D classification yielding final non-uniform refinements (O symmetry) of 2.35 Å for the expanded state (256,929 particles) and 2.50 Å for the contracted state (186,720 particles).

### Cryo-EM image processing of apo-state mjHSP16.5 (75° C)

All steps of cryo-EM image processing were performed in CryoSPARC 4.4.1^62^. The full dataset of 6,460 micrographs for *mj*HSP16.5 apo-75C was preprocessed with Patch Motion Correction (micrographs binned by 2x, 1.0125 Å pixel^-1^) and Patch CTF estimation. The resulting micrographs were culled based on CTF estimation resolution, relative ice thickness, and average intensity to yield 6,186 micrographs carried forward for particle picking. Blob particle picking on the full micrograph stack generated ∼2.4 million picks that were extracted at 2.373 Å pixel^-1^ and cleaned up by two rounds of 2D classification to yield 185,806 particles for further analysis. Results from multi-class *ab initio* generation (4 classes) were input into a heterogeneous refinement (C1 symmetry, 4 classes) which yielded two cage-like maps at 5.69 Å (54,739 particles, set 1) and 4.92 Å (99,715 particles, set 2) resolution. Particles from these two classes were combined and re-extracted at 1.187 Å pixel^-1^ and the pooled particle set (153,807 particles) were reconstructed with O symmetry to 2.86 Å. Results from a C1 consensus non-uniform refinement were input into a 3D Variability analysis with three orthogonal principal modes and a filter resolution of 5 Å^63^.

### Cryo-EM image processing of mjHSP16.5/lysozyme complexes (12:1 ratio)

All steps of cryo-EM image processing were performed in CryoSPARC v4.2.1-4.4.1^62^. The full dataset of 13,276 movies for *mj*:lyso-75C was preprocessed with Patch Motion correction (micrographs binned by 2x, 1.0655 Å pixel^-1^) and Patch CTF estimation. The micrographs were culled based on CTF estimation resolution, relative ice thickness, and average intensity to yield 12,704 micrographs carried forward for particle picking. Blob picking (120–220 Å diameter) on a subset of 500 micrographs to yield 197,163 particles. Particles were extracted at 2.5 Å pixel^-1^, submitted to 2D classification, and the resulting good classes were used as 2D templates for particle picking. Inspection of template-based picks resulted in ∼8.4 million particles that were subjected to two rounds of 2D classification to yield ∼3 million good particles which were then extracted at 3.33 Å pixel^-1^. This particle set was used as input for a multi-class *ab initio* job (8 classes, initial resolution 80 Å). The resulting eight *ab initio* models along with the full good particle stack were input into a heterogeneous refinement job which gave six good classes (2,826,578 total particles) and two noisy classes. A second round of heterogeneous refinement was performed using the six good maps/particles and the two noisy maps (to assist removing noisy particles) which generated four good classes identified as a 24-mer (985,163 particles), 26-mer (450,391 particles), 32-mer (653,689 particles), and a 36-mer oligomeric states (242,970 particles) that were used for further analysis.

The initial particle set pertaining to the 24-mer oligomeric state (985,163 particles) was cleaned up by 2D classification to produce a particle set of 960,040 that was extracted at 1.25 Å pixel^-1^ and input into 3D variability analysis with three orthogonal principal modes and filter resolution of 5 Å^63^. Intermediate mode analysis of the first component was done with five intermediate maps and particles were used as inputs for heterogeneous refinement which produced two classes below 4 Å resolution. These two classes were pooled and refined (non-uniform refinement) without applied symmetry to 2.60 Å resolution.

The particle set associated with the 26-mer oligomeric state class (450,391 particles) was extracted at ∼1.2x binning (1.25 Å pixel^-1^) followed by global CTF refinement and non-uniform refinement without symmetry (C1) to 3.65 Å. 3D variability analysis was performed with three orthogonal principal modes and intermediates analysis of the three components generated five intermediates states that refined to ∼4–8.5 Å resolution without imposed symmetry (C1).

The 32-mer oligomeric state particle set (653,689 particles) was extracted at 1.25 Å pixel^-1^ and cleaned up by 2D classification to give 645,384 particles that refined without applied symmetry to 4.80 Å resolution. 3D variability analysis was performed with three orthogonal principal modes at a filter resolution of 5 Å, followed by intermediates analysis to generate five intermediates states that were used as input for heterogeneous refinement^63^. Heterogeneous refinement produced two classes below 7 Å resolution that corresponded to a 32-mer state (244,887) and a 34-mer state (202,648) that refined (non-uniform refinement) without symmetry (C1) to 4.37 Å and 4.79 Å, respectively. The 34-mer particles went through global CTF refinement and a final non-uniform refinement (C1) to yield a 4.71 Å final map.

The 3x binned particle set for the 36-mer class (242,970) was refined with (D3) and without symmetry (C1) and the resulting maps were used as input for a heterogeneous refinement (C1) with two classes (40 Å initial lowpass filter) and generated one class displaying D3 symmetric features at 8.27 Å resolution with 111,271 particles. Low quality particles were removed by 2D classification resulting in 85,180 particles that refined to 4.50 Å resolution with D3 symmetry. This particle set was expanded with D3 symmetry to yield 511,080 particles that were subjected to local refinement (C1) resulting in a 4.30 Å reconstruction. Output from local refinement was used as input for 3D Variability with three orthogonal principal modes and a filter resolution of 5 Å^63^. Intermediate mode analysis (five intermediates) of the principal components resulted in five intermediate maps with corresponding particle sets that where input for 3D classification without alignment (5 classes). A highly populated class containing 144,177 particles was used for local refinement (C1) resulting in a final reconstruction of the 36-subunit cage structure at 4.01 Å resolution. A reconstruction without symmetry (C1) of the expanded particles was refined to 6.39 Å and used for local resolution analysis. Local resolution estimation of the final refined maps of each oligomeric state was performed in CryoSPARC using default parameters and resolution-based coloring of each map was done in ChimeraX (v1.7)^64^.

### Atomic Modeling, Validation, and Analysis

Atomic model building into the *mj*HSP16.5 apo-state (37° C) in both contracted and expanded states was initiated using a dimer model from the previously published crystal structure of *mj*HSP16.5 (PDB ID: 1SHS^24^). Final maps (O symmetric) corresponding to the contracted and expanded cages were sharpened using Phenix AutoSharpen^65^. Dimers were initially fit as rigid bodies into each map using in ChimeraX to produce a 24-meric model and refinement was done using phenix real space refinement with secondary structure and NCS restraints^66^. Atomic model building of NTD residues 11-32 for the contracted and expanded states were built in COOT as a polyalanine chain, refined, and side chains added before further refinement and side chain adjustment. Iterative manual and automatic model refinement was done in COOT and Phenix (real space refinement) using secondary structure and NCS restraints, and in Isolde without NCS restraints.

The final 24-mer map (C1) from the *mj*HSP16.5/lysozyme (75° C) dataset was sharpened using Phenix AutoSharpen and model building was initiated by rigidly fitting the expanded model from the apo-37C dataset with deletion of residues 11-26^65,66^. Real-space refinement was performed in Phenix using reference model and secondary structure restraints. The final 24-mer model contained residues 27-147. All subsequent model building was initiated using a dimer model from the 24-mer rigidly fit into the final 26-mer, 32-mer, 34-mer, and 36-mer maps. For each oligomeric state various deletions of NTD and CTD residues of the monomers were done to agree with resolved map density and iterative manual remodeling of CTDs to fit the map density was performed in COOT and ISOLDE along with real-space refinement in Phenix^66–68^. Model building of the 26-mer state rigidly fit 13 dimers into the unsharpened map with truncations yielding 22 chains with residues 34-147 and 4 chains with residues 34-139. Sixteen dimers were refined in the sharpened (Phenix local anisotropic sharpening) 32-mer map, resulting in 31 chains with residues 31-147 and one chain with residues 35-143. For the 34-mer state, 17 dimers refined into a Phenix auto-sharpened map, resulting in 33 chains covering residues 33-147 and one chain with residues 35-143. The 36-mer state was built with 18 refined dimers fit into a Phenix AutoSharpened map with 36 chains covering residues 31-147. For all models, validation of model refinement and map-to-model fit was done using Phenix validation and the PDB validation server^65^.

For visualization of unmodeled density, final maps were low-pass filtered at 7 Å resolution and density corresponding to the atomic models of each state were generated using the molmap (7 Å) function in ChimeraX v1.17.1. Density corresponding to the 7 Å molmaps were subtracted from the respective 7 Å low-pass filtered Cryo-EM maps to generate a subtracted map containing the unmodeled internal density. Cα r.m.s.d. calculations were generated in ChimeraX using only chains with full CTDs (through residue 147) resulting in 24 chains for the 24mer, 22 chains for the 26mer, 31 chains for the 32mer, 33 chains for the 34mer, and 36 chains for the 36mer for this comparative analysis. Coloring based on Cα r.m.s.d. and local resolution was done in ChimeraX with the color by attributes and surface color utilities, respectively. Buried surface areas for the ACD-dimer interface, inter-dimer interface, and the canonical CTD/ACD-groove interface were calculated using the Interfaces function in ChimeraX with default settings except areaCutoff set to 100 Å^2^. For visualization, modeling of NTD residues 1-10 was (Fig 2e) was done by extension of residues distally from residue 11 of the contracted model and subsequent addition of side chains and refinement in COOT. Measurement of the internal volume density (at ∼2*σ*) was performed in ChimeraX (v1.7) by subtracting cage density using molmaps generated at 7 Å and the volume subtraction tool. PDB 1DPX was used for measurement of lysozyme volume. Assessment of non-NTD internal density volume was performed by calculating the volume of residues 1-32 from a 7 Å molmap, multiplying by the number of subunits to determine the total volume occupied by NTDs, and subtracting this from the total internal volume density of the cryo-EM map (Supplemental Movie 4).

### Figure Preparation

Structural models and cryo-EM density maps were visualized and prepared for presentation using ChimeraX. Final figures were composed in Photoshop.

### AI-assisted technologies

During the preparation of this work the authors used ChatGPT to help revise portions of the text to improve readability. After using this tool, the authors reviewed and edited the content as needed and take full responsibility for the content of the publication.

## DATA AVAILABILITY

Cryo-EM density maps have been deposited to the Electron Microscopy Data Bank (EMD-XXXXX). Coordinates for atomic models have been deposited to the Protein Data Bank (XXXX). The original multi-frame micrographs have been deposited to EMPIAR (EMPIAR-XXXXX). Plasmids used for protein expression are available upon request. Raw data from DLS and NS-EM are provided on zenodo doi: xxxx.

## EXTENDED DATA (Tables, Figure and Legends)

**Extended Data Table 1.**
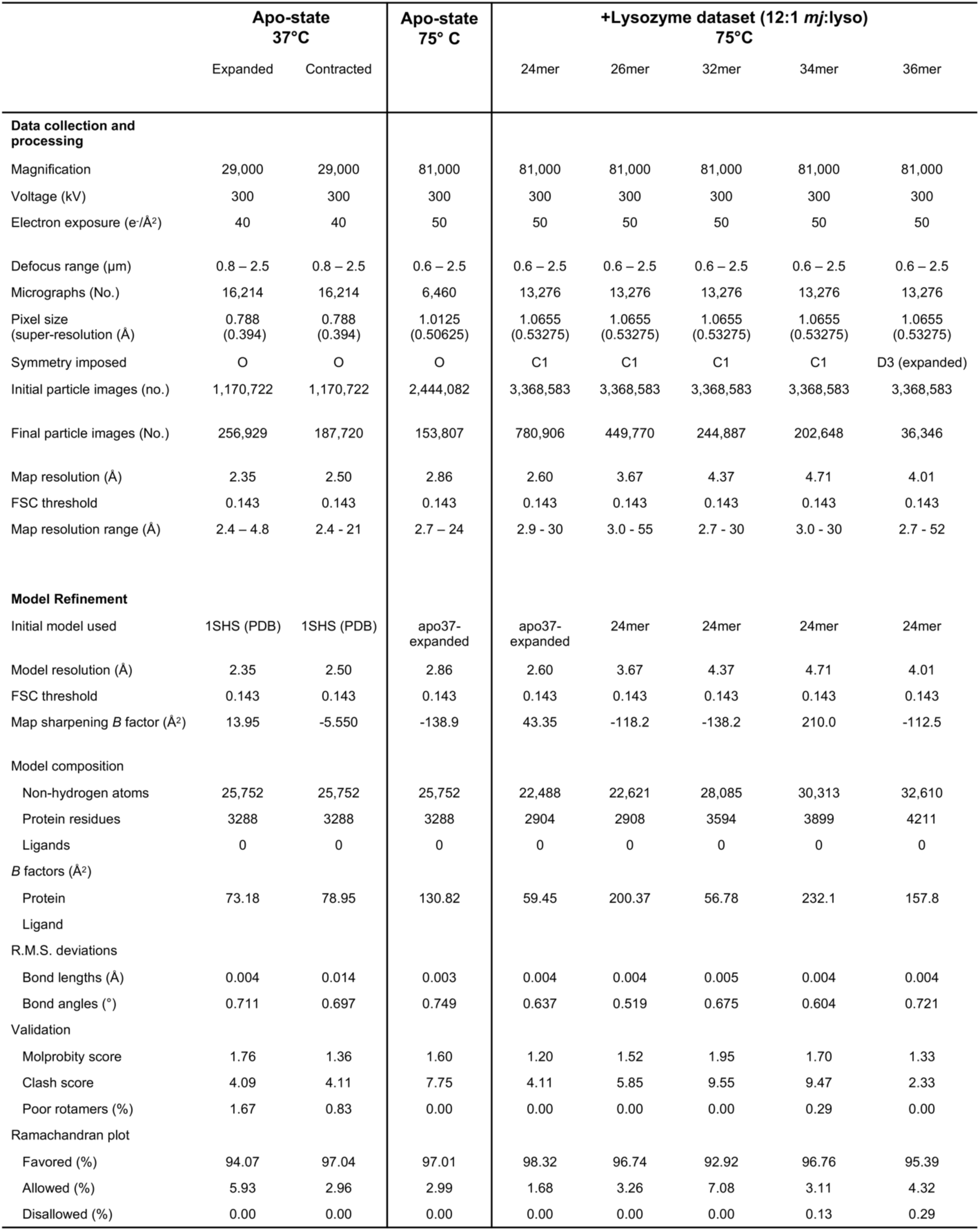
Summary of single-particle cryo-EM data collection, processing, model building, and validation for maps and models.

|  | Apo-state<br>37°C |  | Apo-state<br>75° C | +Lysozyme dataset (12:1 *mj:lyso*)<br>75°C |  |  |  |  |
| --- | --- | --- | --- | --- | --- | --- | --- | --- |
|  | Expanded | Contracted |  | 24mer | 26mer | 32mer | 34mer | 36mer |
| Data collection and processing |  |  |  |  |  |  |  |  |
| Magnification | 29,000 | 29,000 | 81,000 | 81,000 | 81,000 | 81,000 | 81,000 | 81,000 |
| Voltage (kV) | 300 | 300 | 300 | 300 | 300 | 300 | 300 | 300 |
| Electron exposure (e-/Å²) | 40 | 40 | 50 | 50 | 50 | 50 | 50 | 50 |
| Defocus range (µm) | 0.8 – 2.5 | 0.8 – 2.5 | 0.6 – 2.5 | 0.6 – 2.5 | 0.6 – 2.5 | 0.6 – 2.5 | 0.6 – 2.5 | 0.6 – 2.5 |
| Micrographs (No.) | 16,214 | 16,214 | 6,460 | 13,276 | 13,276 | 13,276 | 13,276 | 13,276 |
| Pixel size<br>(super-resolution (Å)) | 0.788<br>(0.394) | 0.788<br>(0.394) | 1.0125<br>(0.50625) | 1.0655<br>(0.53275) | 1.0655<br>(0.53275) | 1.0655<br>(0.53275) | 1.0655<br>(0.53275) | 1.0655<br>(0.53275) |
| Symmetry imposed | O | O | O | C1 | C1 | C1 | C1 | D3 (expanded) |
| Initial particle images (no.) | 1,170,722 | 1,170,722 | 2,444,082 | 3,368,583 | 3,368,583 | 3,368,583 | 3,368,583 | 3,368,583 |
| Final particle images (No.) | 256,929 | 187,720 | 153,807 | 780,906 | 449,770 | 244,887 | 202,648 | 36,346 |
| Map resolution (Å) | 2.35 | 2.50 | 2.86 | 2.60 | 3.67 | 4.37 | 4.71 | 4.01 |
| FSC threshold | 0.143 | 0.143 | 0.143 | 0.143 | 0.143 | 0.143 | 0.143 | 0.143 |
| Map resolution range (Å) | 2.4 – 4.8 | 2.4 - 21 | 2.7 – 24 | 2.9 - 30 | 3.0 - 55 | 2.7 - 30 | 3.0 - 30 | 2.7 - 52 |
| Model Refinement |  |  |  |  |  |  |  |  |
| Initial model used | 1SHS (PDB) | 1SHS (PDB) | apo37-expanded | apo37-expanded | 24mer | 24mer | 24mer | 24mer |
| Model resolution (Å) | 2.35 | 2.50 | 2.86 | 2.60 | 3.67 | 4.37 | 4.71 | 4.01 |
| FSC threshold | 0.143 | 0.143 | 0.143 | 0.143 | 0.143 | 0.143 | 0.143 | 0.143 |
| Map sharpening *B* factor (Å²) | 13.95 | -5.550 | -138.9 | 43.35 | -118.2 | -138.2 | 210.0 | -112.5 |
| Model composition |  |  |  |  |  |  |  |  |
| Non-hydrogen atoms | 25,752 | 25,752 | 25,752 | 22,488 | 22,621 | 28,085 | 30,313 | 32,610 |
| Protein residues | 3288 | 3288 | 3288 | 2904 | 2908 | 3594 | 3899 | 4211 |
| Ligands | 0 | 0 | 0 | 0 | 0 | 0 | 0 | 0 |
| *B* factors (Å²) |  |  |  |  |  |  |  |  |
| Protein | 73.18 | 78.95 | 130.82 | 59.45 | 200.37 | 56.78 | 232.1 | 157.8 |
| Ligand |  |  |  |  |  |  |  |  |
| R.M.S. deviations |  |  |  |  |  |  |  |  |
| Bond lengths (Å) | 0.004 | 0.014 | 0.003 | 0.004 | 0.004 | 0.005 | 0.004 | 0.004 |
| Bond angles (°) | 0.711 | 0.697 | 0.749 | 0.637 | 0.519 | 0.675 | 0.604 | 0.721 |
| Validation |  |  |  |  |  |  |  |  |
| Molprobity score | 1.76 | 1.36 | 1.60 | 1.20 | 1.52 | 1.95 | 1.70 | 1.33 |
| Clash score | 4.09 | 4.11 | 7.75 | 4.11 | 5.85 | 9.55 | 9.47 | 2.33 |
| Poor rotamers (%) | 1.67 | 0.83 | 0.00 | 0.00 | 0.00 | 0.00 | 0.29 | 0.00 |
| Ramachandran plot |  |  |  |  |  |  |  |  |
| Favored (%) | 94.07 | 97.04 | 97.01 | 98.32 | 96.74 | 92.92 | 96.76 | 95.39 |
| Allowed (%) | 5.93 | 2.96 | 2.99 | 1.68 | 3.26 | 7.08 | 3.11 | 4.32 |
| Disallowed (%) | 0.00 | 0.00 | 0.00 | 0.00 | 0.00 | 0.00 | 0.13 | 0.29 |

**Extended Data Table 2.**
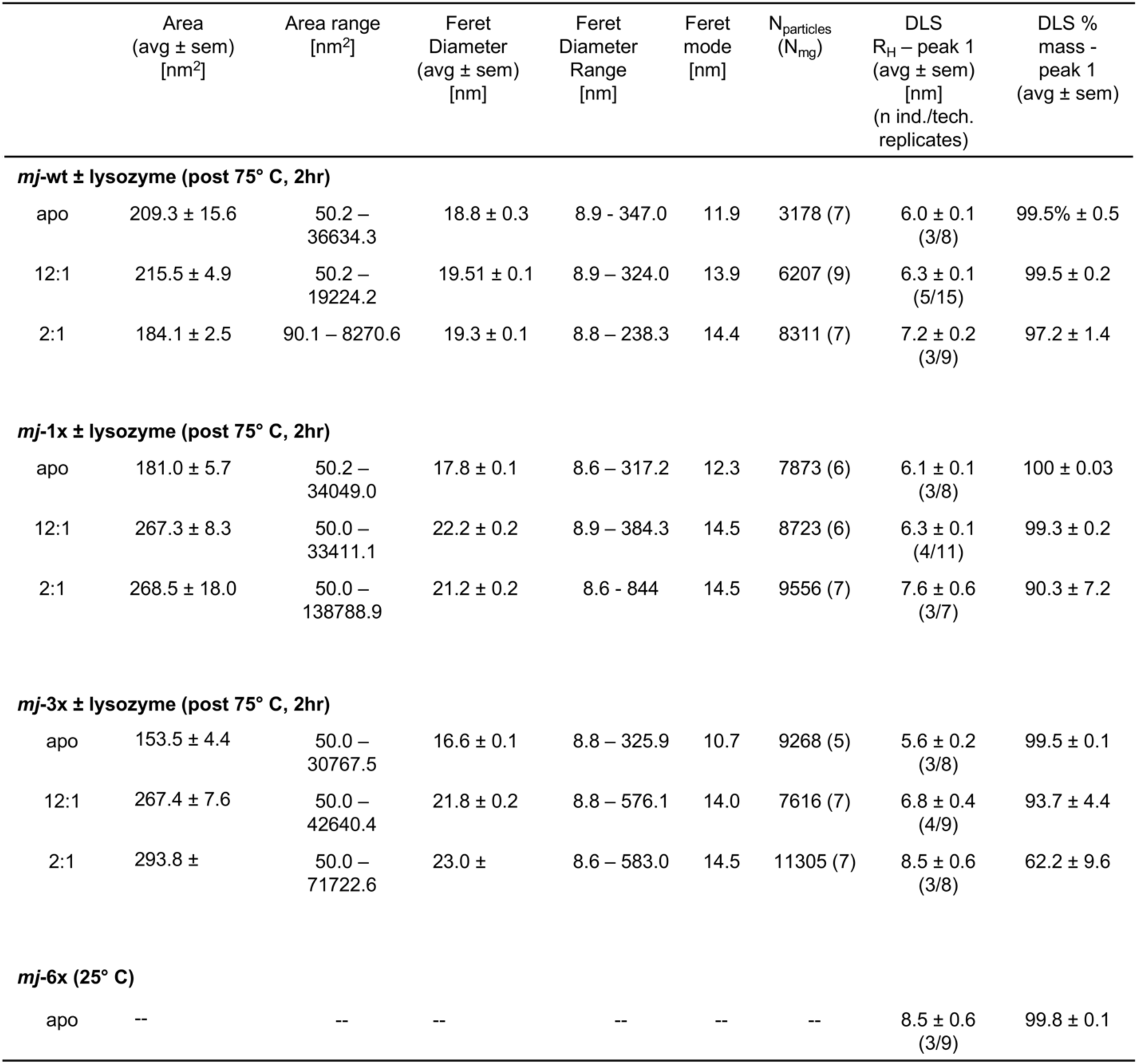
Single-particle analysis from the NS-EM datasets (Area and Feret diameter) and DLS measurements (R_h_; hydrodynamic radius and corresponding percent mass) of *mj*HSP16.5 wildtype and NTD variants obtained in the presence and absence of client lysozyme. NS-EM grids were prepared following two hour incubation at 75° C. The exception is for the *mj*-6x variant which was done at 25° C due to reduced thermal stability and measurements obtained by hand (n=100). Measurements are presented as average ± s.e.m.

**Extended Data Figure 1.**
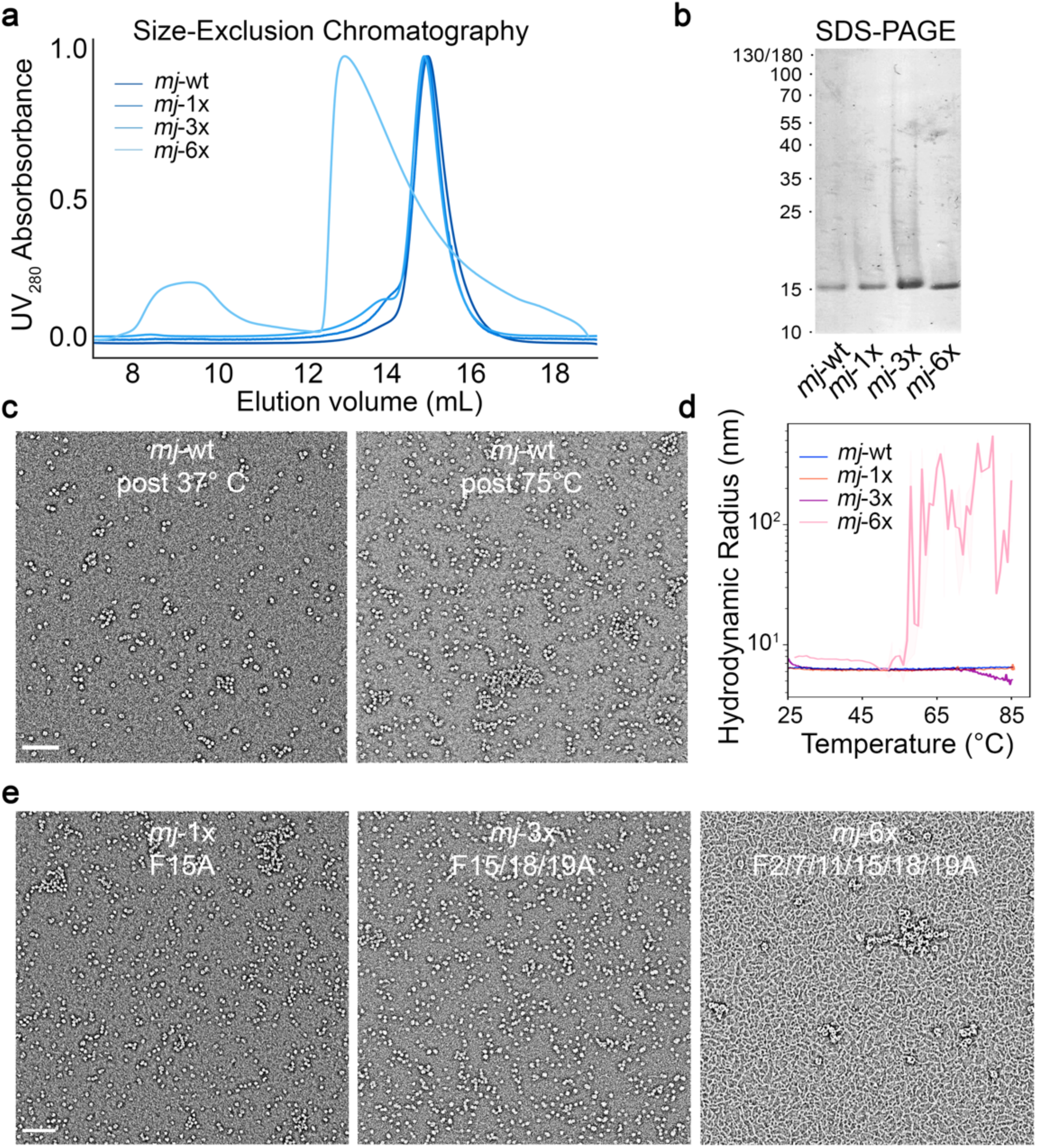
Purification, negative-stain EM and thermos-stability of *mj*HSP16.5 constructs. **a**, Size-exclusion chromatography traces for apo-states of *mj*HSP16.5 wildtype (*mj*-wt) and NTD variants (*mj*-1x, *mj*-3x, and *mj*-6x). **b**, SDS-PAGE visualized by silver staining of purified *mj*-wt, *mj*-1x, *mj*-3x, and *mj*-6x. **c**, Negative-stain electron microscopy (NS-EM) of apo-state *mj*HSP16.5 after heating at 37° C (∼16 hours) and 75° C (2 hours). **d**, Temperature ramp from 25-85° C for *mj*-1x, *mj*-3x, and *mj*-6x and corresponding hydrodynamic radius showing *mj*-6x aggregation around 55-60° C. *mj*-wt and other variants were stable up to 85° C. **e**, NS-EM of apo-state *mj*-1x (75° C) and *mj*-3x (75° C) and *mj*-6x (25° C) *mj*-6x at two dilutions displayed background fibers (∼2 nm wide) and circular assemblies. Micrograph scale bars = 100 nm.

**Extended Data Figure 2.**
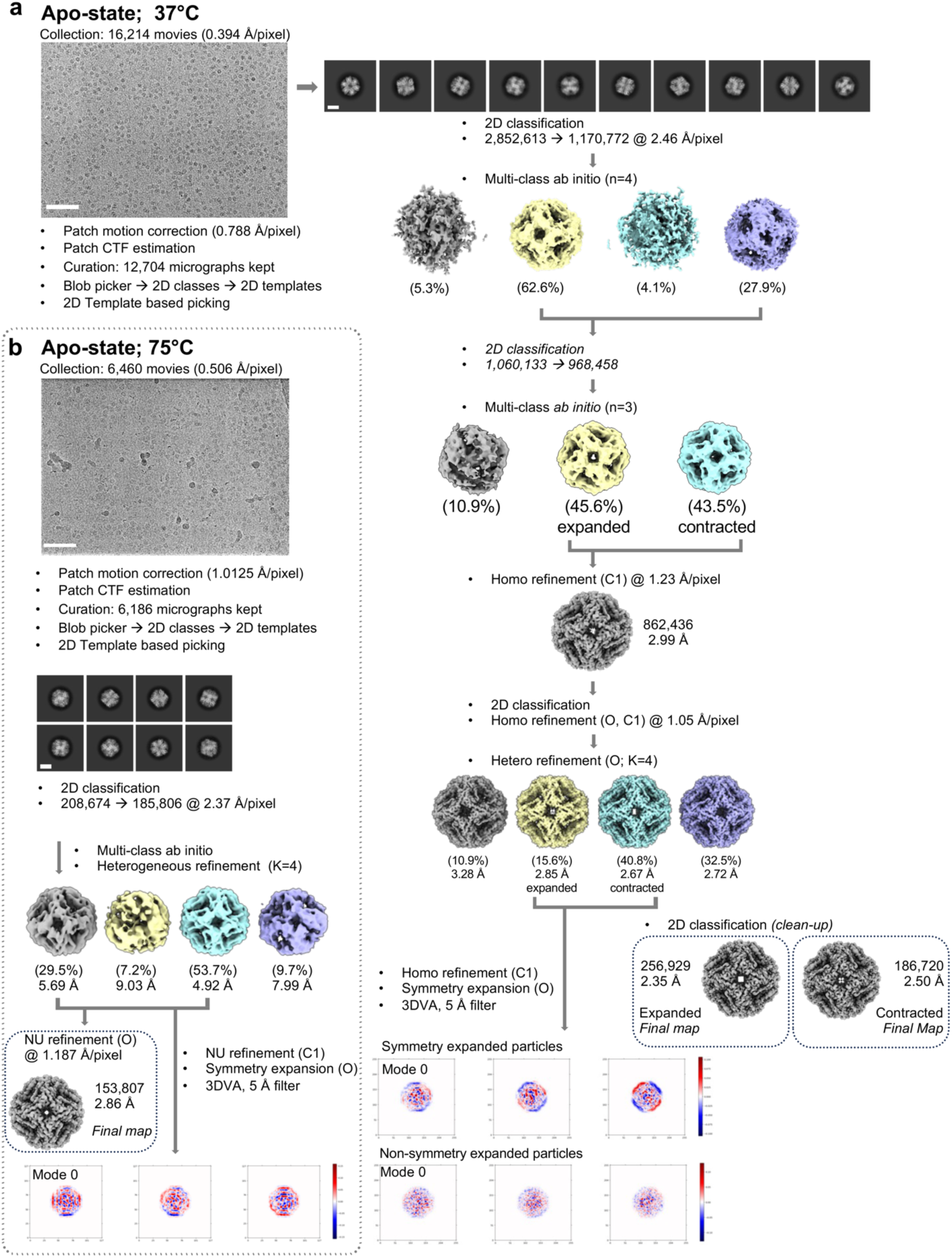
Single-particle cryo-EM image processing workflows for the *mj*HSP16.5 apo-37 and apo-75 datasets. Overview of preprocessing, 2D/3D classification, *ab initio* model generation, 3D refinement, and 3D variability analysis steps for a, apo-state 37° and b, apo-state 75° datasets. Particle count numbers, pixel sizes, symmetries, and resolutions are noted where appropriate. Scale bars for micrographs = 100 nm, and for 2D classes = 5 nm.

**Extended Data Figure 3.**
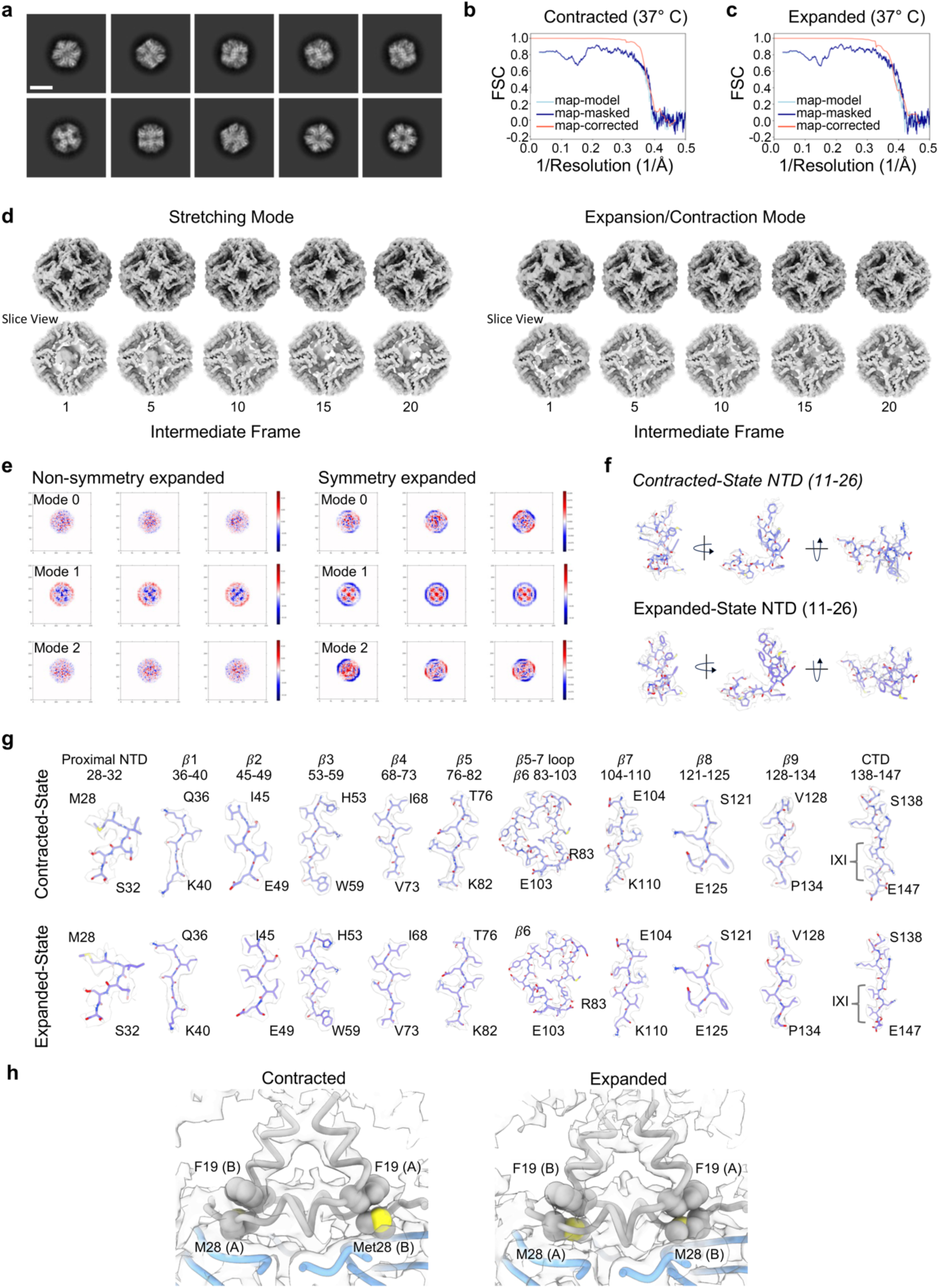
Resolution assessment and 3D variability analysis of the apo-37 cryo-EM dataset. **a**, Representative 2D classes showing multiple views of the canonical 24-mer caged assembly. Scale bar = 10 nm. **b-c**, Fourier Shell Correlation (FSC) plots of the contracted and expanded states displaying the CryoSPARC generated FSC plot of the corrected map (red), the unmasked map to model FSC (light blue) and masked map to model FSC (dark blue) generated by Phenix. **d**, Intermediate reconstructions for the stretching and expansion principal component modes identified from 3D variability analysis in CryoSPARC, shown along 4-fold axis (top) and internal slice view (bottom). **e**, Principal component modes from 3D variability analysis with and without symmetry expansion of the consensus particle set, displayed as heat maps of density variability across each mode. **f-g**, Segmented views of the model-to-map fit for the contracted and expanded atomic models. **h**, Interaction of Phe19 and Met28 (shown as space-filling model) between dimeric protomers (A and B).

**Extended Data Figure 4.**
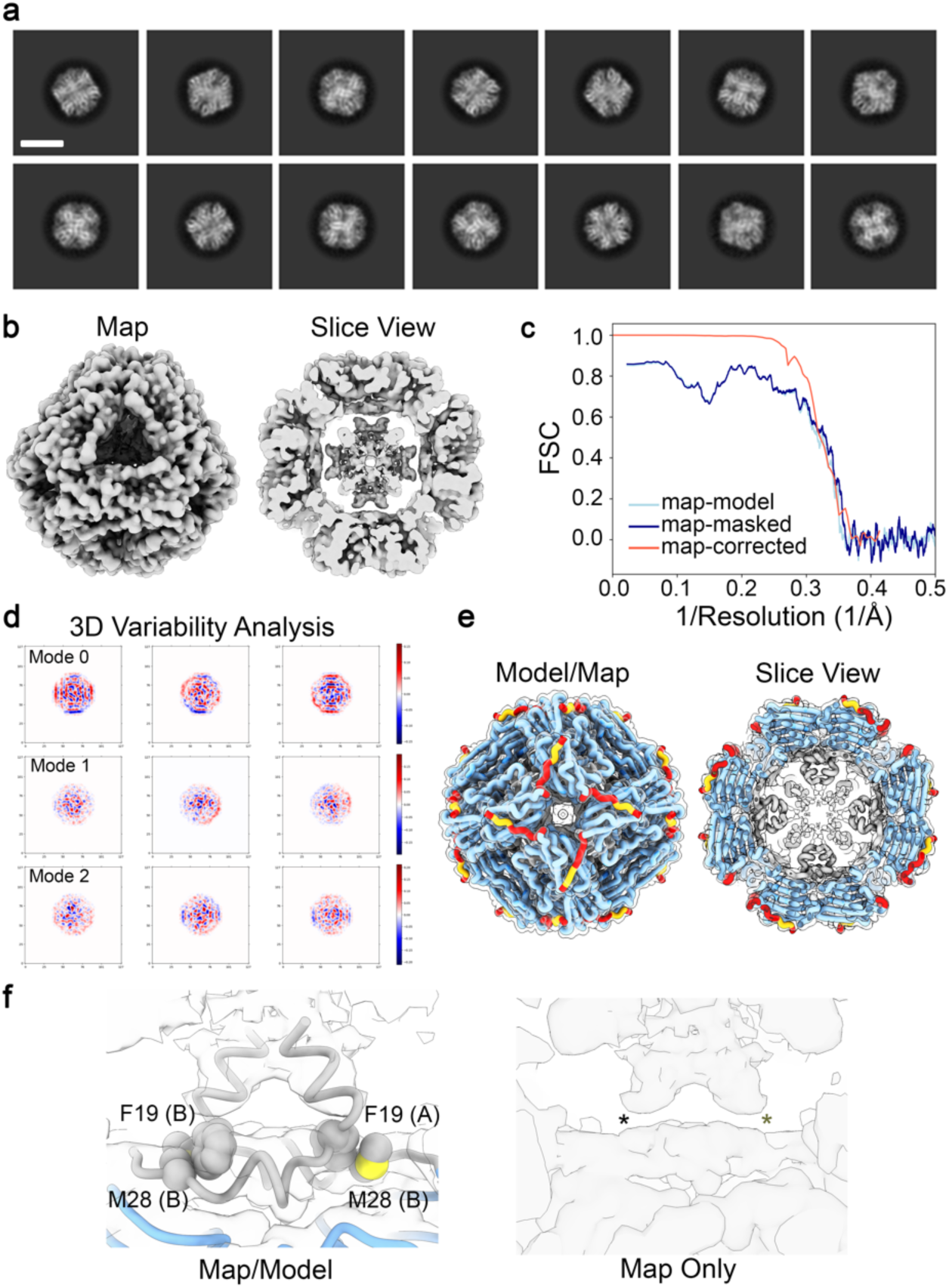
Resolution assessment and 3D variability analysis of the apo-75 cryo-EM dataset. **a**, Representative 2D classes showing multiple views of the canonical 24mer caged assembly. Scale bar = 10 nm. **b**, Consensus 3D reconstruction of apo-75 with octahedral (O) symmetry imposed, shown from the canonical 3-fold axis of the 24-mer (left) and slice view displaying internal density (right) showing helical density from the NTD protruding toward the center of the cage. **c**, Fourier shell correlation (FSC) plot of apo-75 reconstruction show in (b), displayed for the corrected map (red) from CryoSPARC, the unmasked map to model FSC (light blue) and masked map to model FSC (dark blue) from Phenix. **d**, Principal component modes from 3D variability analysis displayed as heat maps of density variability across each mode. **e**, Atomic model for residues 11-147 fit into sharpened cryo-EM density map. **f**, *Left*, Map and model showing putative competition between Phe18 and Phe19 with Met28 between dimeric protomer chains (A and B). *Right*, Map without model showing weakened density at base of NTD helix *α*1, as compared to the apo-37 maps.

**Extended Data Figure 5.**
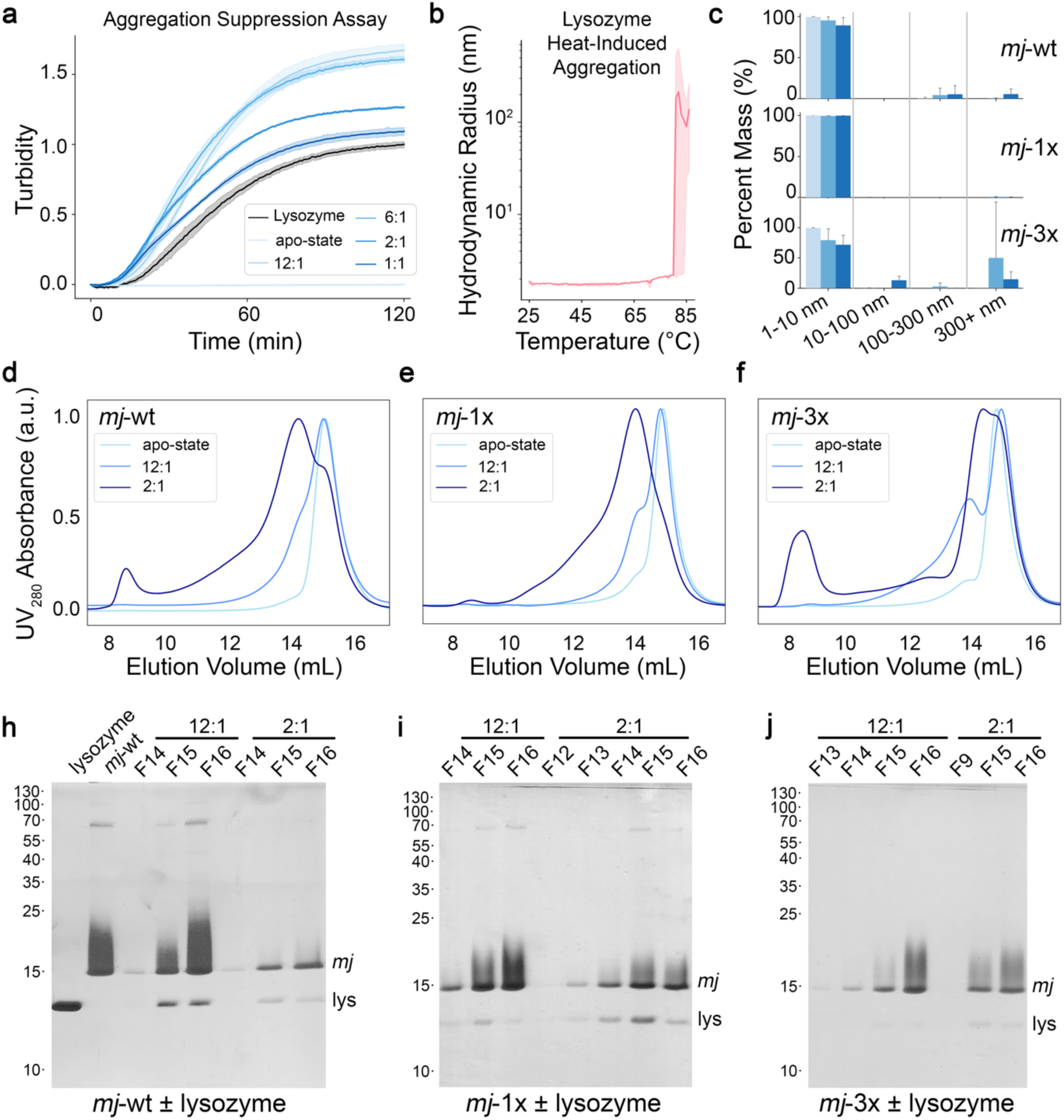
Chaperone-client binding assays for *mj*HSP16.5 wildtype (*mj*-wt) and NTD variants (*mj*-1x: F15A; *mj*-3x: F15/18/19A). **a,** Overlay of turbidity traces obtained at 37° C for ratios of *mj*-wt to lysozyme (12:1, 6:1, 2:1, and 1:1), *mj*-wt alone (light blue), and lysozyme alone (gray). For these conditions, lysozyme unfolding was initiated with reducing agent. Trace shows average and s.d. of n=4 technical replicates. b, Bulk hydrodynamic radius of lysozyme, under non-reducing conditions, measured from 25°–85° C by dynamic light scattering showing heat-induced aggregation occurring at temperatures above ∼80° C. Trace shows average and s.d. of n=3 replicates. **c,** Histograms showing binned hydrodynamic radii (1-10 nm, 10-100 nm, 100-300 nm, and 300+ nm bins) and associated percent mass for *mj*-wt, *mj*-1x, and *mj*-3x in the presence and absence of lysozyme, under non-reducing conditions, after incubation at 75° C for two hours. Error bars represent 95% confidence interval; n = 3-5 independent experiments. d-f, Representative size-exclusion chromatography (SEC) traces for *mj*-wt (d), *mj*-1x (e), and *mj*-3x (f) with lysozyme (12:1 and 2:1 chaperone:client ratios) and in the apo-state after incubation at 75° C for two hours. **h-j**, SDS-PAGE analysis of fractions collected from SEC runs of *mj*-wt, *mj*-1x, and *mj*-3x with lysozyme (12:1 and 2:1 ratios), respectively. Position of molecular weight markers indicated (left) and protein bands corresponding the *mj*HSP16.5 (*mj*) and lysozyme (lys) indicated (right).

**Extended Data Figure 6.**
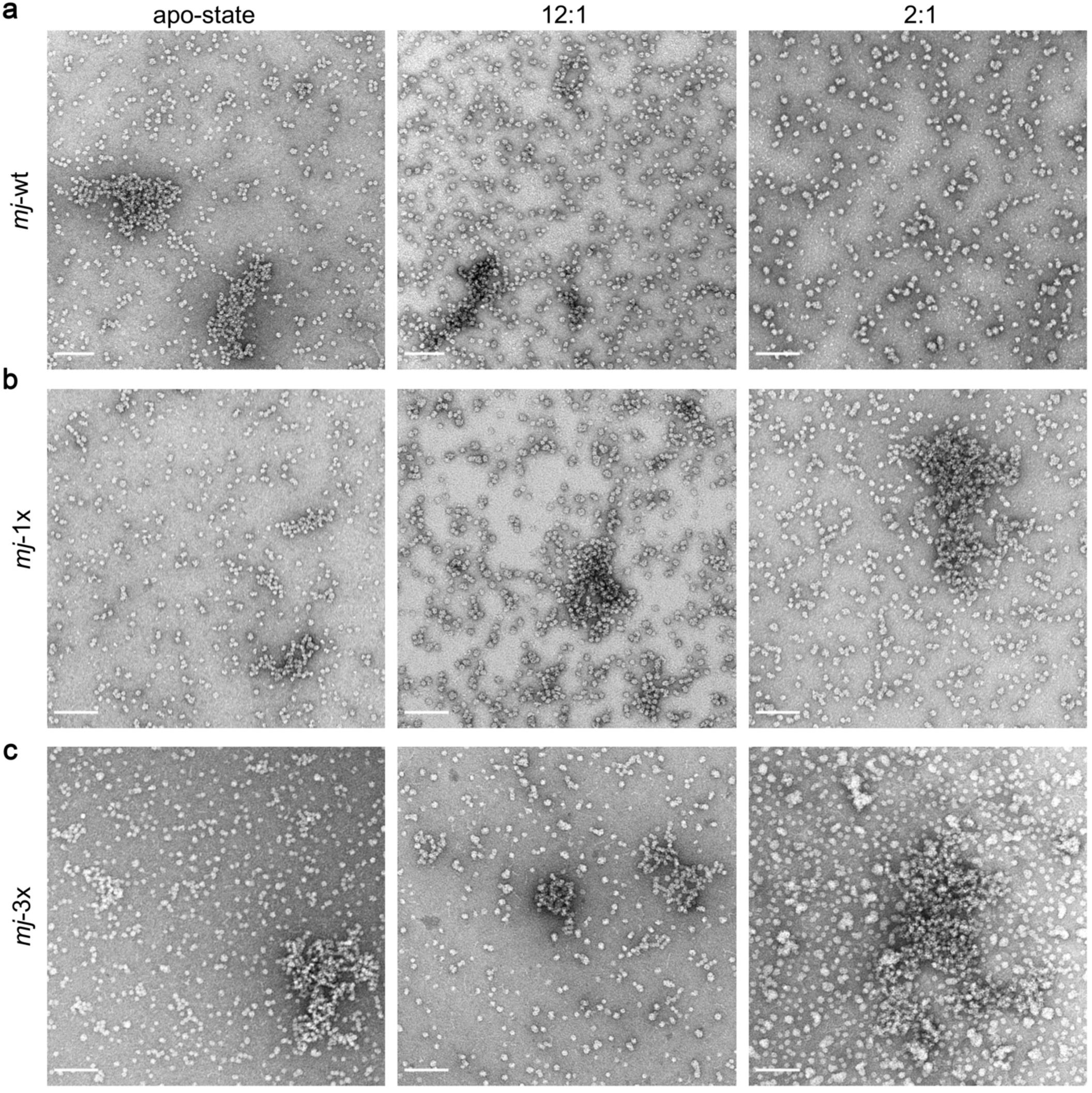
NS-EM analysis of lysozyme chaperone assays. Representative unfiltered NS-EM micrographs of a, *mj*-wt, b, *mj*-1x, and c, *mj*-3x in the apo-state (*left*), and for the 12:1 (*middle*) and 2:1 (*right*) chaperone:client ratios. Scale bar = 100 nm.

**Extended Data Figure 7.**
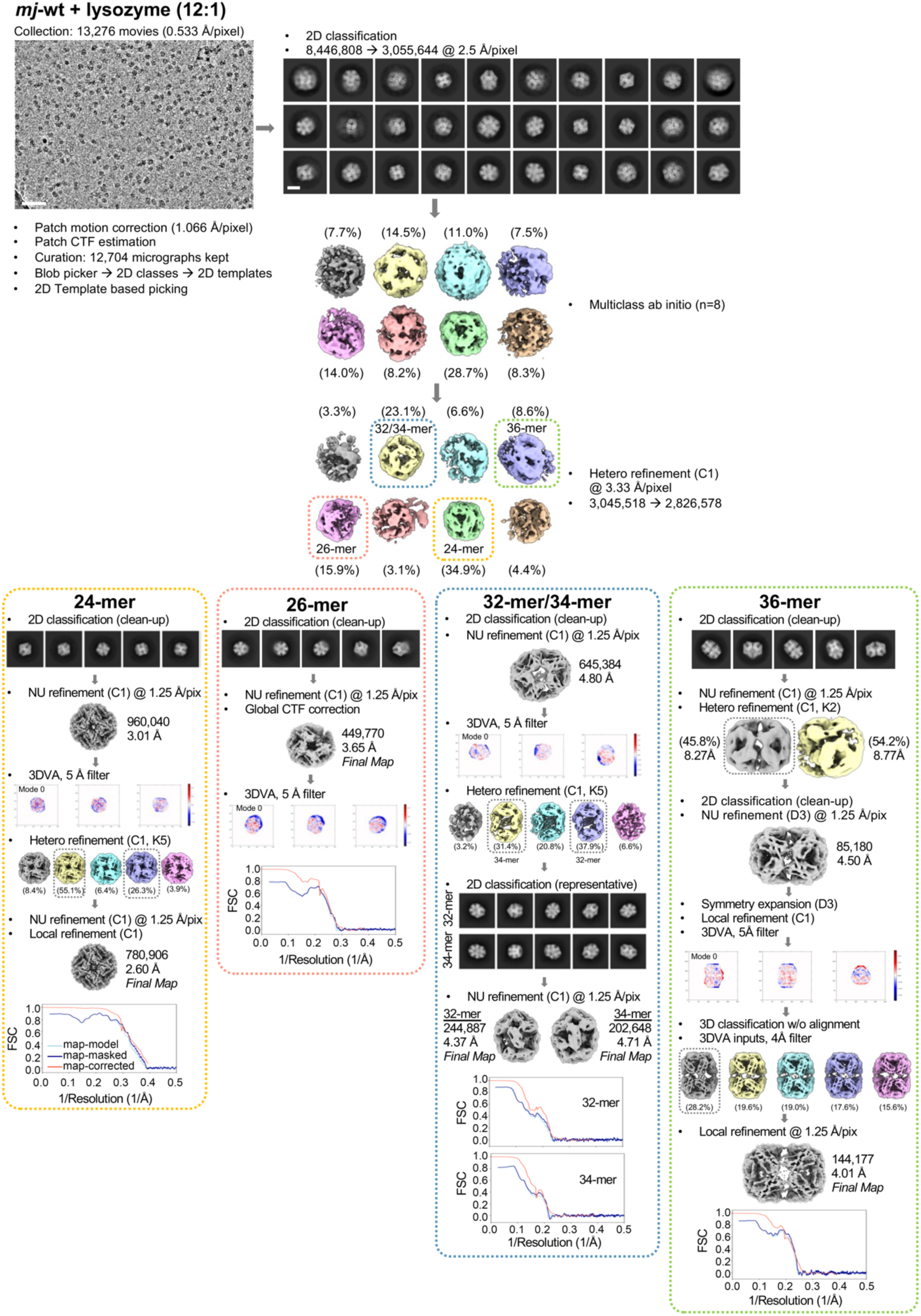
Single-particle cryo-EM processing workflow for the *mj*HSP16.5/lysozyme dataset. Overview of preprocessing, 2D/3D classification, *ab initio* model generation, 3D refinement, and 3D variability analysis steps. Particle count numbers, pixel sizes, symmetries, and resolutions are noted were appropriate. Outlined boxes are color coded to match *ab initio* models and corresponding downstream processing for each oligomeric state. Final maps used for model building are noted along with the corresponding CryoSPARC generated Fourier Shell Correlation (FSC) plot for the corrected map (red), and unmasked map to model FSC (light blue) and masked map to model FSC (dark blue) generated by Phenix. Scale bars for micrograph = 100 nm, and for 2D classes = 10 nm.

